# CDK1 and CDK2 regulate phosphorylation-dependent NICD1 turnover and the periodicity of the segmentation clock

**DOI:** 10.1101/245704

**Authors:** Francesca Anna Carrieri, Philip Murray, Paul Davies, Jacqueline Kim Dale

**Affiliations:** Division of Cell and Developmental Biology, School of Life Sciences, University of Dundee, Dow Street, Dundee, DD15EH, Scotland, UK; Department of Mathematics, University of Dundee, Nethergate, Dundee, DD14HN, Scotland, UK; MRC Phosphorylation Unit, School of Life Sciences, University of Dundee, Dow Street, Dundee, DD15EH, Scotland, UK

## Abstract

All vertebrates share a segmented body axis. Segments form periodically from the rostral end of the presomitic mesoderm (PSM) and this periodicity is regulated by the segmentation clock, a molecular oscillator that drives dynamic clock gene expression across the PSM with a periodicity that matches somite formation. Notch signalling is crucial to this process. Altering Notch intracellular domain (NICD) stability affects both the clock period and somite size. However, the mechanistic details of how NICD stability is regulated are unclear.

We identified a highly conserved site crucial for NICD recognition by the SCF E3 ligase, which targets NICD for degradation. We demonstrate both CDK1 and CDK2 can phosphorylate NICD in the domain where this crucial residue lies and that NICD levels vary in a cell cycle-dependent manner. Inhibiting CDK1 or CDK2 activity increases NICD levels both *in vitro* and *in vivo*, leading to a delay of clock gene oscillations.

## INTRODUCTION

Segmentation, a process which occurs early during vertebrate body plan formation, generates repeated segments (or somites) that later give rise to the vertebral column, most skeletal musculature and dermis [1, 2].

During somitogenesis, pairs of somites bud off the rostral end of the unsegmented presomitic mesoderm (PSM) with a periodicity that is species specific. The periodicity of segment formation is regulated by a molecular oscillator, known as the somitogenesis clock, which drives oscillatory gene expression within the PSM tissue from which somites are derived [2–4].

These clock genes are targets of the Notch, Wnt and FGF pathways [5, 6]. Aberrant somitogenesis leads to severe segmentation and skeletal defects [7]. In humans, defects in segmentation lead to congenital scoliosis (CS), with an infant mortality rate of 50% that comprises many vertebral skeletal and muscular pathologies, including the family of spondylocostal dysostoses (SCD). For CS, whilst the aetiology is unclear, linkage analyses have shown mutations in four genes lead to familial forms of SCD [8]. Significantly these are components of the Notch pathway, which plays multiple roles during segmentation. Notch is crucial to the segmentation process in mice, since in the absence of Notch signalling, the segmentation clock stops and no somites form [9].

On a single cell level in the PSM, oscillatory clock gene expression is established through positive and negative feedback loops of unstable clock gene products which potentiate or inhibit the pathway that activates them. Synchronisation of clock gene oscillations between neighbouring cells is reliant on Notch signalling [10–13]. Mathematical models predict the period of clock gene oscillations can be approximated as a sum of the delays involved in transcription, splicing, translation and transport of clock gene products, and in particular through the regulation of the half-lives of both mRNA and protein of unstable regulators [14–17]. Whilst great progress has been made in demonstrating the role of transcription and splicing delays in setting the clock period, little experimental work investigating whether stability of clock components affects clock period has been performed.

Most studies addressing the molecular mechanisms regulating the periodicity of clock gene oscillations have focused on the role of Notch signalling components [10, 18–23] Notch is one of the major highly conserved signalling pathways that regulate cell-cell communication which involves gene regulation mechanisms that control multiple processes during development and adult life [24–31].

Upon extracellular ligand activation, Notch transmembrane receptors are cleaved, releasing the intracellular domain (NICD) that translocates to the nucleus to regulate expression of specific developmental gene cohorts [30, 32, 33]. NICD is highly labile, and phosphorylation-dependent turnover acts to restrict Notch signalling [34–36].

All known canonical Notch activity relies on this regulation of NICD half-life. Moreover, aberrant NICD turnover contributes to numerous cancers and diseases [24, 28, 37–42]. Despite the multiple impacts of NICD turnover in both development and disease, the molecular details regulating this turnover remain uncharacterised. The stability of NICD and therefore duration of the Notch signal is regulated by phosphorylation of the C-Terminal PEST domain which leads to subsequent recruitment of FBXW7, F-Box and WD Repeat Domain Containing 7, (a key component of the SCF^Sel10/FBXW7^ E3 ubiquitin ligase complex) [34–36, 43–49]. Ultimately, this leads NICD to ubiquitylation and proteasomal degradation [43, 50–53]. However, the molecular details of NICD degradation mediated by FBXW7 are not well understood.

A recent study combining experimental and computational biology demonstrated changes in NICD stability affect the chick and mouse somitogenesis clock period which in turn affects somite size. In this study a pharmacological approach was used to demonstrate that culturing chick/mouse PSM explants with broad specificity inhibitors of cyclin-dependent protein kinases (Roscovitine/DRB) and Wnt signalling (XAV939) leads to elevated levels and a prolonged NICD half-life and phase shifted clock oscillation patterns both at a tissue level and in larger segments. Furthermore, reducing NICD production in this assay rescues these effects [18]. These results imply potential coupling between NICD degradation and the segmentation clock. However, the specific kinases/molecular mechanism of action remain ill-defined and leave open the question of whether this coupling is a general or conserved mechanism.

In this manuscript we identify the phosphorylated residues within human NICD. We demonstrate that purified recombinant Cyclin-dependent kinase 1 (CDK1) and Cyclin-dependent kinase 2 (CDK2) phosphorylate NICD within the PEST domain. A point mutation affecting a conserved serine residue within this CDK substrate domain of the NICD PEST motif prevents NICD interaction with endogenous FBXW7. Strikingly, we show that NICD levels fluctuate in a cell cycle dependent manner anti-correlating with high levels of CDK1/2 activity. Lastly, highly specific inhibitors of CDK1 or CDK2 lead to increased levels of NICD *in vitro* and *in viv*o and delay the mouse somitogenesis clock and somite formation.

Using a mathematical model we show that the experimental observations made in cell lines and PSM tissue can be explained in a single theoretical framework that couples the cell cycle to NICD degradation.

## RESULTS

### Roscovitine, DRB and XAV939 increase NICD levels in HEK293, iPS and IMR90 cells

Direct phosphorylation of NICD in its PEST domain enhances its turnover and thus degradation [34–36]. A study using broad range kinase inhibitors demonstrated that the stability and turnover of NICD is linked to the regulation of the pace of the segmentation clock across the PSM in chick and mice embryos [18]. However, this study did not define the specific kinases or molecular mechanism of action of the inhibitors. In order to identify which kinases are involved in NICD phosphorylation and which residues in the NICD PEST domain are phosphorylated rendering NICD susceptible to degradation, we employed a cellular model due to the limiting quantity of material available using embryonic cell lysates.

First, we used the same inhibitors as Wiedermann *et al*. [18], and thus investigated if Roscovitine, DRB and XAV939 elicit the same effect upon NICD levels in a variety of cell culture models, namely HEK293 (human embryonic kidney), iPS (induced pluripotent stem cells) and IMR90 (human Caucasian foetal lung fibroblast) cells.

Roscovitine is a small molecule belonging to the family of purines. It inhibits cyclin-dependent kinases (CDKs) through direct competition with ATP for binding at the ATP-binding site of CDKs [54, 55]. DRB (5,6-dichloro-1-β-D-ribofuranosylbenzimidazole) also inhibits CDKs, particularly CDK7 and 9 [56, 57]. XAV939 is a Wnt inhibitor that stimulates β-catenin degradation by stabilizing axin through inhibition of the enzymes tankyrase 1 and 2 [58]. HEK293, iPS or IMR90 cells treated for 3 hours with each of the three inhibitors at the same concentrations used in embryonic lysate studies [18] led to an increase in NICD levels compared to control cells cultured in the presence of DMSO (**Figures 1A-D, Supplementary Figure 1**). In control conditions, NICD was not easily detectable due to its very short half-life. Quantification of the density of western blot bands in at least three independent experiments confirmed that the increase in NICD levels was statistically significant after treatment with Roscovitine, DRB and XAV939, as shown in **Figures 1B, 1D and Supplementary Figure 1B**. Two other inhibitors were used as positive and negative controls for the assay. LY411575 is a γ-secretase inhibitor that prevents Notch1 cleavage and thus inhibits activation of target gene expression [59, 60]. As expected, LY411575 treatment significantly reduced NICD levels (**Figures 1A-D, Supplementary Figure 1**). Phosphorylation of the C-Terminal PEST domain of NICD leads to recruitment of FBXW7 and thus to NICD ubiquitylation and proteasomal degradation [34–36, 43, 44, 46–48]. When E3 ligase activity is reduced with the NEDDylation inhibitor MLN4924 [61], NICD levels increase, since NICD degradation is stopped in the presence of this compound (**Figures 1A-D, Supplementary Figure 1**).

**Figure 1.**
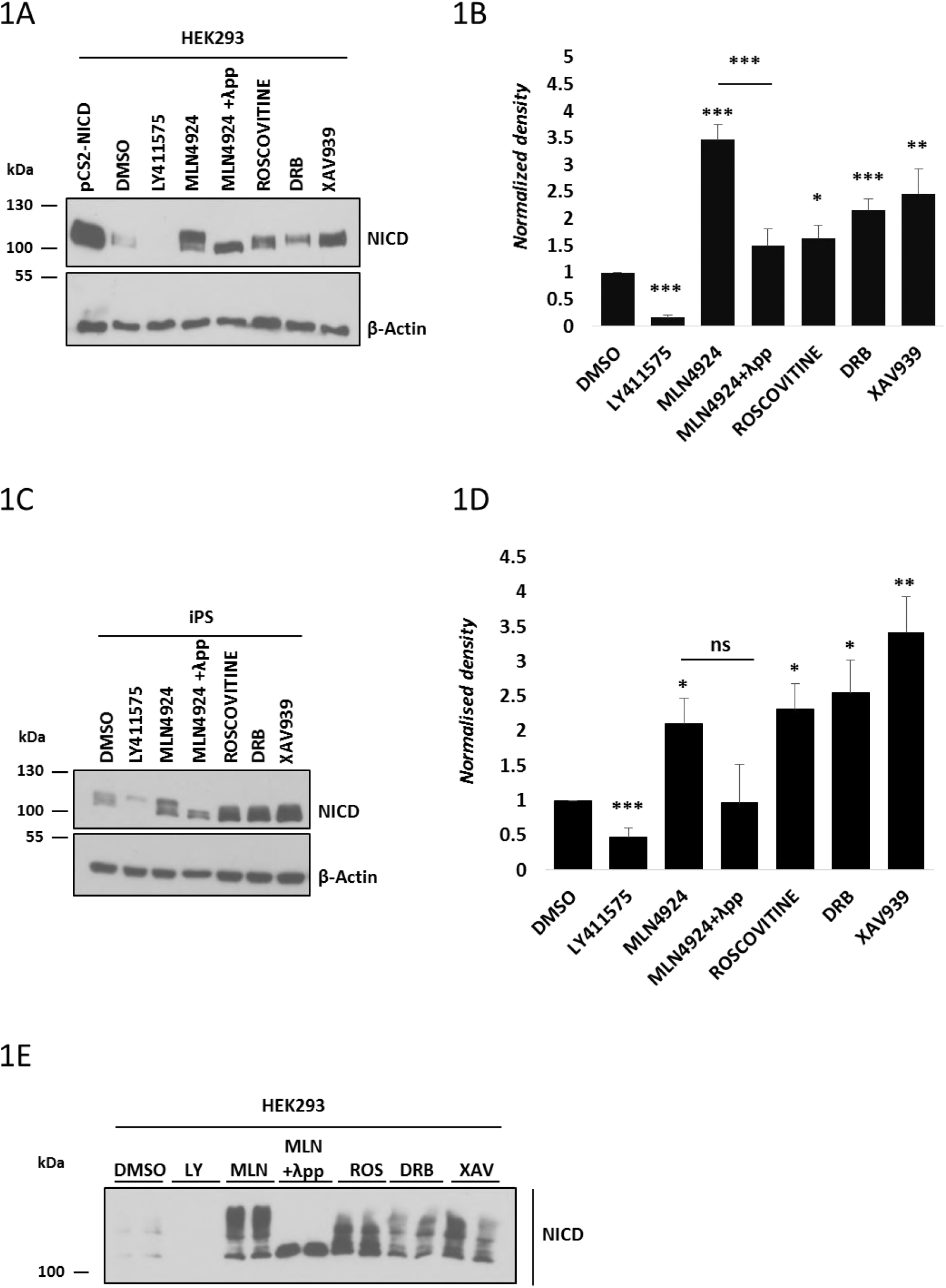
Endogenous NICD levels increase in HEK293 cells following treatment with Roscovitine, DRB and XAV939. (A) HEK293 cells were treated for 3 hours with 150 nM of LY411575, 1 μM of MLN4924, 10 μM of Roscovitine, 10 μM DRB or 10 μM XAV939. DMSO served as vehicle control. Transfection with pCS2-NICD vector served as positive control. Western blot analysis reveals that NICD levels were increased upon treatment with Roscovitine, DRB, XAV939, or MLN4924. NICD is undetectable following LY411575 treatment. NICD antibody detects a doublet following inhibitor treatment and the top band disappears following λ phosphatase treatment indicating that this top band is a phosphorylated isoform. β-Actin served as loading control. (B) Quantification of the density of western blot bands in (A) using ImageJ software. Data are expressed as fold changes compared to DMSO treatment. All data represent the mean ± SEM from three independent experiments. Student’s t-test analysis was performed with *p≤0.05, **p≤0.01, ***p≤0.001. (C) iPS cells were treated for 3 hours with 150 nM of LY411575, 1 μM of MLN4924, 10 μM of Roscovitine, 10 μM DRB or 10 μM XAV939. DMSO has been used as vehicle control. Western blot analysis reveals that NICD levels were increased upon treatment with Roscovitine, DRB, XAV939, or MLN4924. NICD is undetectable following LY411575 treatment. NICD antibody detects a doublet following Inhibitor treatment and the top band disappears following λ phosphatase treatment indicative that this top band is a phosphorylated isoform. β-Actin served as loading control. (D) Quantification of the density of western blot bands in (C) using ImageJ software. Data are expressed as fold changes compared to DMSO treatment. All data represent the mean ± SEM from three independent experiments. Student’s t-test analysis was performed with *p≤0.05, **p≤0.01, ***p≤0.001. (E) HEK293 cells were treated with the same inhibitors as described in (A). NICD phosphorylation status was analysed by a Phos-tag assay. DMSO served as vehicle control. NICD phosphorylation profile varies following Roscovitine, DRB, XAV939 or MLN4924 treatment. Following λ phosphatase treatment none of the high molecular weight bands are visible indicating they are all phosphorylated isoforms.

Interestingly, we were able to detect two distinct bands by western blot with the NICD antibody, particularly when cells were treated with MLN4924. We hypothesised this reflected the presence of non-phospho and phospho-NICD species. To test this hypothesis, we treated MLN4924-treated lysates with λ phosphatase which abrogated the appearance of the higher band by western blot with the NICD antibody (**Figures 1A and 1C**). These data demonstrate that the higher band detected corresponds to a phosphorylated isoform of NICD.

In order to determine if the increased levels of NICD were due to increased NICD production and/or increased NICD stability, we exposed HEK293 cells to LY411575 treatment for the last hour of culture, thereby inhibiting the production of new NICD. Under control conditions with LY411575 treatment in the last hour, NICD levels are very low (**Supplementary Figure 2, lane 2**). However, despite LY411575 treatment in the last hour, cells cultured in the presence of small molecule inhibitors showed increased levels of NICD compared to the control, indicating that the increase in NICD levels is not due to increased NICD production, but to an increased stability (**Supplementary Figure 2**).

Taken together, these results show that exposure to this group of inhibitors leads to increased levels of NICD in a variety of cell lines, in the same way that they do in the mouse and chicken PSM tissue, suggesting they regulate a conserved mechanism leading to increased NICD levels and reduced NICD turnover.

### Phostag analysis following Roscovitine, DRB and XAV939 treatment reveals a variety of NICD phospho-species

In order to investigate whether this selection of small molecule inhibitors have different effects on NICD phosphorylation, we treated HEK293 cells with the inhibitors and performed a Phostag assay [62]. Following MLN4924 treatment, a variety of bands indicative of different phospho-species of NICD was observed (**Figure 1E**). Given that very few bands are present in the control sample (DMSO, **Figure 1E**), the bands detected after MLN4924 treatment are likely to be very labile isoforms of the NICD peptide, which are rapidly degraded in the DMSO sample. In contrast, when whole cell lysate was treated with both MLN4924 and λ phosphatase only one band, of the lowest molecular weight, was detectable, further supporting the notion that the ladder of bands obtained upon MLN4924 treatment reflects a variety of unstable phosphorylated NICD isoforms, which is completely depleted in the presence of λ phosphatase (**Figure 1E**). As expected from data showing NICD levels are increased after inhibitor treatment (**Figures 1A-D**), all three kinase inhibitors Roscovitine, DRB and XAV939 cause a noticeable increase in the number and intensity of bands compared to control cells. However, compared to the effect seen with MLN4924, phos-tag technology reveals Roscovitine, DRB and XAV939 have a reduced number of phospho-bands as compared to MLN4924, indicative of the fact these kinase inhibitors act to reduce NICD phosphorylation. Moreover, each of the inhibitors presents a distinct profile of NICD phospho-species. These data suggest that NICD is targeted by several kinases and/or phosphorylation events which are differentially sensitive to these inhibitors (**Figure 1E**).

NICD-FBXW7 interact at the endogenous levels in HEK293 cells, in a phosphorylation-dependent manner. The involvement of the F-box protein component of the SCF E3 ligase complex, FBXW7, in NICD degradation has been previously reported [46–48]. However, to date, the NICD-FBXW7 interaction has only been shown in overexpressed systems [35, 44, 46, 63–66] Thus, we examined the binding of NICD to FBXW7 using co-immunoprecipitation analysis at the endogenous level.

FBXW7 was immunoprecipitated from HEK293 cells treated with DMSO or MLN4924 for 3 hours and extracts were probed with NICD antibody. NICD directly binds to FBXW7 (**Figure 2A**). After MLN4924 treatment, the amount of NICD bound to FBXW7 was significantly higher compared to control cells and this was particularly evident with the higher molecular weight isoform of NICD (**Figures 2A-B**), confirming again NICD-FBXW7 interaction is phosphorylation-dependent.

**Figure 2.**
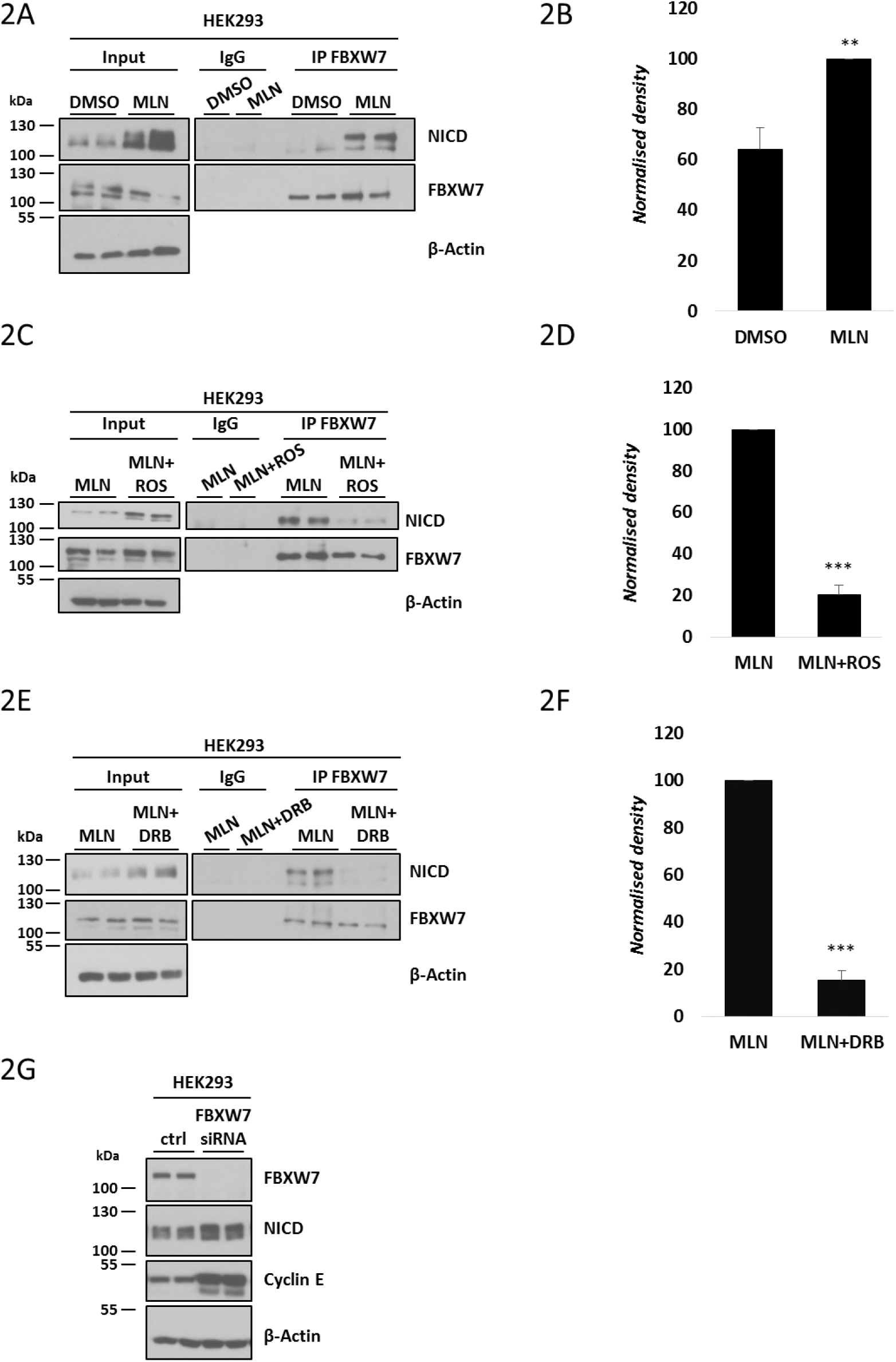
NICD and FBXW7 interact directly at endogenous levels. (A) NICD interaction with FBXW7 at endogenous levels in HEK293 cells. 500 μg of HEK293 cell lysates treated with DMSO or MLN4924 were subjected to immunoprecipitation using FBXW7 antibody, or IgG antibody as negative control, and precipitated material was analysed by western blot using NICD antibody. Western blot with FBXW7 antibody served as loading control for immunoprecipitation efficiency. 10% of cell lysate before immunoprecipitation was used as input control and β-Actin served as loading control. (B) Quantification of the density of western blot bands in (A) performed by ImageJ software. Data are expressed as fold changes compared to DMSO. All data represent the mean ± SEM from three independent experiments. Student t-test was used to determine p values, with **p≤0.01. (C) Roscovitine treatment reduced the NICD-FBXW7 interaction. 500 μg of HEK293 cell lysates treated with MLN4924 or MLN4924 together with Roscovitine were subjected to immunoprecipitation using FBXW7 antibody, or IgG antibody as negative control, and precipitated material was analysed by western blot using NICD antibody. Western blot with FBXW7 antibody served as loading control for immunoprecipitation efficiency. 10% of cell lysate before immunoprecipitation was used as input control and β-Actin served as loading control. (D) Quantification of the density of western blot bands in (C) performed by ImageJ software. Data are expressed as fold changes compared to MLN4924 treated samples. All data represent the mean ± SEM from three independent experiments. Student’s t-test analysis was performed, with ***p≤0.001. (E) Interaction between NICD and FBXW7 is reduced following DRB treatment. 500 μg of HEK293 cell lysates treated with MLN4924 or MLN4924 together with DRB were subjected to immunoprecipitation using FBXW7 antibody, or IgG antibody as negative control, and precipitated material was analysed by western blot using NICD antibody. Western blot with FBXW7 antibody served as loading control for immunoprecipitation efficiency. 10% of cell lysate before immunoprecipitation was used as input control and β-Actin served as loading control. (F) Quantification of the density of western blot bands in (E) performed by ImageJ software. Data are expressed as fold changes compared to MLN4924 treated samples. All data represent the mean ± SEM from three independent experiments. Student’s t-test analysis was performed, with ***p≤0.001. (G) Inhibiting endogenous FBXW7 modulates endogenous NICD levels. HEK293 cells were transfected with control (scrambled siRNA) or FBXW7 siRNA. Levels of FBXW7, NICD and Cyclin E were determined by western blot. β-Actin served as loading control.

In order to determine if the change in the NICD phosphorylation profile observed after treatment with the CDK inhibitors reduced the NICD-FBXW7 interaction, we performed the same co-immunoprecipitation assay after CDK inhibitor treatment. In order to maximise the amount of NICD immunoprecipitated with FBXW7, cells were treated with MLN4924 (to prevent NICD degradation) in the presence or absence of the CDK inhibitors. A significantly reduced interaction between NICD and FBXW7 was observed after treating HEK293 cells with Roscovitine or DRB for 3 hours (**Figures 2C and 2E**). Statistical analyses, carried out on the density of western blot bands after immunoprecipitation, confirmed a significant reduction in the interaction between NICD and FBXW7 following either Roscovitine or DRB treatment (**Figures 2D and 2F**).

Taken together, these data demonstrate, for the first time, that NICD interacts with FBXW7 at endogenous levels in HEK293 cells, and this interaction is dependent on phosphorylation. To further validate the involvement of FBXW7 in endogenous NICD turnover, we conducted a siRNA-mediated depletion of FBXW7 in HEK293 cells (**Figure 2G**). siRNA treatment efficiently depleted FBXW7 protein levels and led to an increase in levels of the FBXW7 target protein Cyclin E. FBXW7 depletion also resulted in increased levels of NICD, and, in particular, an accumulation of the phosphorylated form of NICD (**Figure 2G**).

### Serine 2513 is essential for the NICD-FBXW7 interaction

We utilised Mass Spectrometry as an unbiased approach to identify which NICD residues are phosphorylated in HEK293 cells transiently transfected with human NICD-GFP, followed by immunoprecipitation of NICD-GFP. Gel slices were processed and submitted to MS analysis. We identified 15 phospho-sites on exogenous hNICD, highlighted in green in **Figure 3A**. To investigate the relevance of those phosphorylation sites in NICD turnover, we screened those located within the PEST domain (such as S2527), and others based on the FBXW7 phosphodegron motif (such as S2205, S2513, S2516, S2538) which is known to be [RK] S/T P [RK] X S/T/E/D, where X is any amino acid and RK is any amino acid except arginine (R) or lysine (K) [67].

**Figure 3.**
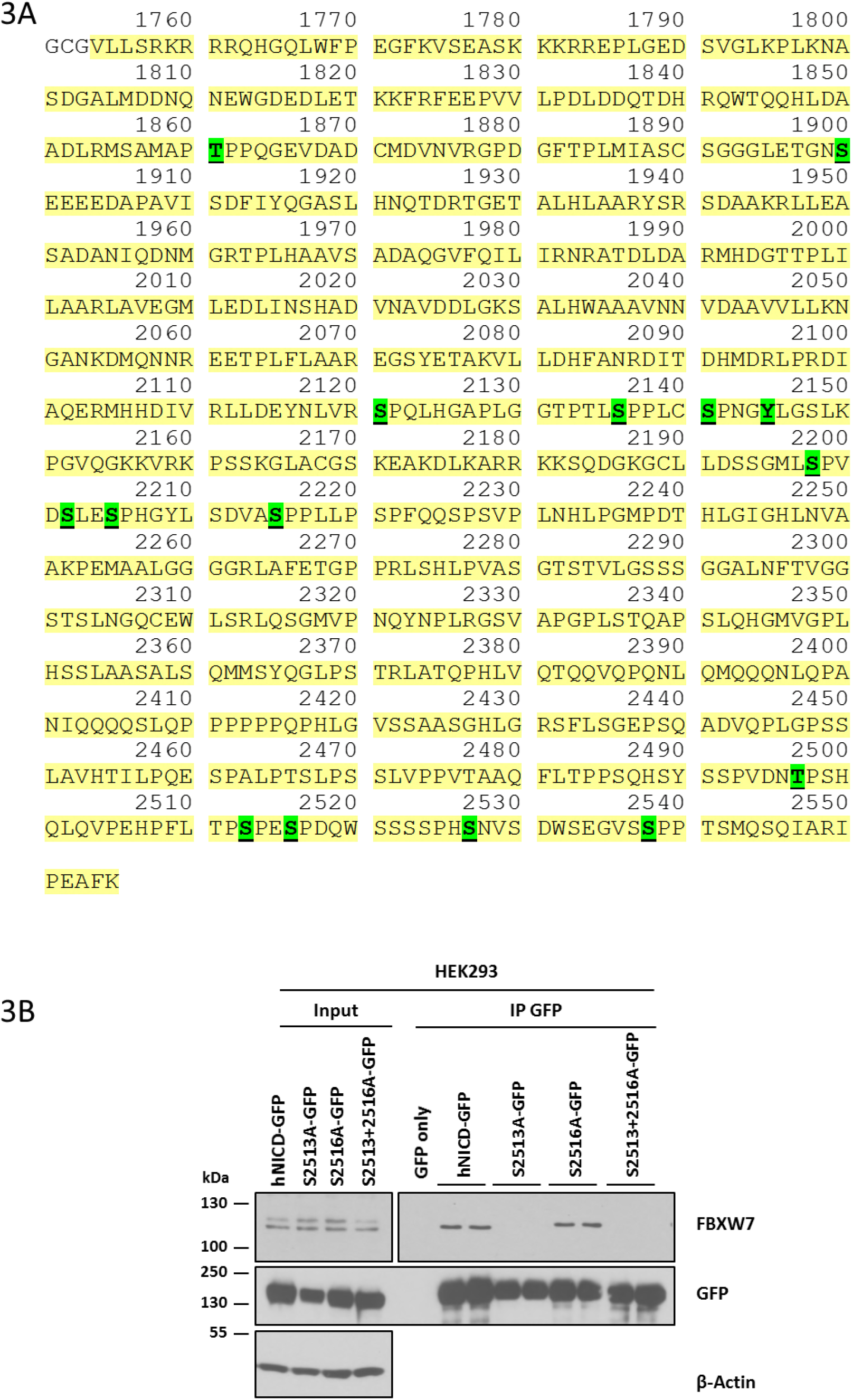
Mass spectrometry analysis of phosphorylated residues in hNICD. (A) LC-MS-MS analysis of in-gel-digested HEK293 cells transfected with hNICD-GFP, and subjected to immunoprecipitation using GFP antibody, identified multiple phosphorylation sites in NICD, highlighted in green. (B) Phosphorylation of serine 2513, but not serine 2516, is required for the NICD-FBXW7 interaction. hNICD-GFP phospho-mutant peptides encoding non-phosphorylatable residues at S2513 and/or 2516 (serine to alanine) were expressed in HEK293 cells. The exogenously expressed protein was subsequently immunoprecipitated with anti-GFP antibody and precipitated material was analysed by western blot using FBXW7 antibody. Wild-type hNICD-GFP and GFP only vectors were included as positive and negative controls, respectively. Western blot using GFP antibody served as immunoprecipitation efficiency control. β-Actin has been used as loading control for the input lanes.

Thus, we generated 5 peptides each carrying a serine to alanine point mutation in an identified site. Following transient transfection of HEK293 cells with wild-type or mutated peptides we performed immunoprecipitation using GFP-conjugated beads, to evaluate peptide binding efficiency with endogenous FBXW7.

In 2004, Fryer and colleagues proposed that CDK8 phosphorylates serine residues 2481, 2484, 2506 (2513, 2516, 2538 in our annotation). They reported that when all of these residues were mutated from serine to alanine, *in vitro* phosphorylation by recombinant CyClinC:CDK8 was dramatically reduced [35], suggesting these sites could be implicated in NICD stability. However, we find individual mutations on serine residues 2205, 2516, 2527 and 2538 did not affect the NICD-FBXW7 interaction (**Figure 3B, Supplementary Figure 3**). However, mutating serine 2513 to alanine, to render this residue non-phosphorylatable, completely abolished the NICD-FBXW7 interaction (**Figure 3B**). Cells transfected with the double mutant S2513A/S2516A also showed a dramatic loss of the NICD-FBXW7 interaction. This did not reflect a reduction in the level of immunoprecipitated GFP (**Figure 3B**).

Thus, our data suggest that only serine 2513, of those we have tested, is the key NICD phosphorylation site required for interaction with FBXW7, and thus potentially crucial for NICD stability and turnover.

### Cyclin Dependent Kinase (CDK) 1 and 2 phosphorylate NICD *in vitro*

Previous reports have proposed a number of potential kinases which may be involved in NICD phosphorylation and turnover, including CyclinC:CDK8 [35], CyclinC:CDKs [65] and GSK3β [34, 36]. The MRC-PPU Kinase Profiling Inhibitor Database (http://www.kinasescreen.mrc.ac.uk/kinase-inhibitors) (University of Dundee) indicates that, at the concentrations used in our assays, Roscovitine, in particular, is a potent inhibitor of CDK2, but a very weak inhibitor of CK1, GSK3β and more than 50 other kinases tested. In order to confirm results derived from the Kinase Profiling Inhibitor Database, we performed a kinase assay in collaboration with the MRC-PPU International Centre for Kinase Profiling. We tested three NICD phospho-peptides against the activity of a panel of different kinases including CDK1 and CDK2 (**Table 1**). Of seven kinases tested, 5 had no specific activity against any of the peptides. In contrast, CDK1 and CDK2 elicited a very high activity against Peptide 1, which contained serine residues 2513 and 2516, previously identified by Mass spectrometry analysis to be phosphorylated in NICD and in particular 2513 we have shown to be crucial for the NICD-FBXW7 interaction.

**Table 1.** CDK1 and CDK2 exhibit specific activity against a NICD phospho-peptide by *in vitro* kinase assay. Three NICD phospho-peptides were tested for the activity of seven different kinases (CDK1, CDK2, CDK5, CDK7, CDK9, CK1α and GSKβ). A known substrate was used as control for each kinase. The specific activity of each kinase is expressed in *U/mg*. Peptide 1 contains serine residues 2513 and 2516 (HPFLTP<u>**S**</u>PE<u>**S**</u>PDQWSSSSPH). Peptide 2 includes serine 2538 (NVSDWSEGVS**<u>S</u>**PPTSMQSQIA). Peptide 3 encompasses serine residue 2141 (GTPTLSPPLC**<u>S</u>**PNGYLGSLKP).

| *Specific activity (U/mg)* | *Peptide 1* | *Peptide 2* | *Peptide 3* | *Control* |
| --- | --- | --- | --- | --- |
| Cdk1/CyclinA2 | 1035.43 | 35.21 | -7.86 | 1322 |
| Cdk1/CyclinB | 887.23 | 37.46 | 0.04 | 1566 |
| Cdk2/CyclinA | 79.7 | 0.5 | 0 | 117.7 |
| Cdk5/p35 | 47 | 0.8 | 0.5 | 1729.5 |
| Cdk7/MAT1/CyclinH | 0.1 | 0.2 | 0.1 | 21.6 |
| Cdk9/CyclinT1 | 2.7 | 0.6 | 0.3 | 66.4 |
| CK1α | 0 | 0 | 0.1 | 19.1 |
| GSKβ | 1 | 1 | 1.1 | 552.5 |

**Peptide 1: S2513 and S2516** HPFLTPSPESPDQWSSSSPH **Peptide 2: S2538** NVSDWSEGVSPPPTSMQSQIA **Peptide 3: S2141** GTPTLSPPLCSPNGYLGSLKP

These results demonstrate that CDK1 and CDK2 phosphorylate the C-terminal region of NICD *in vitro*. Therefore, we decided to evaluate the contribution of these kinases to endogenous NICD turnover and FBXW7 interaction in the cell lines. CDK2 siRNA treatment in HEK293 cells efficiently depleted CDK2 protein levels, with no effect on levels of CDK4. Under these conditions NICD levels were significantly increased compared to control scrambled siRNA treated cells, indicative of reduced NICD turnover (**Figures 4A-B**).

**Figure 4.**
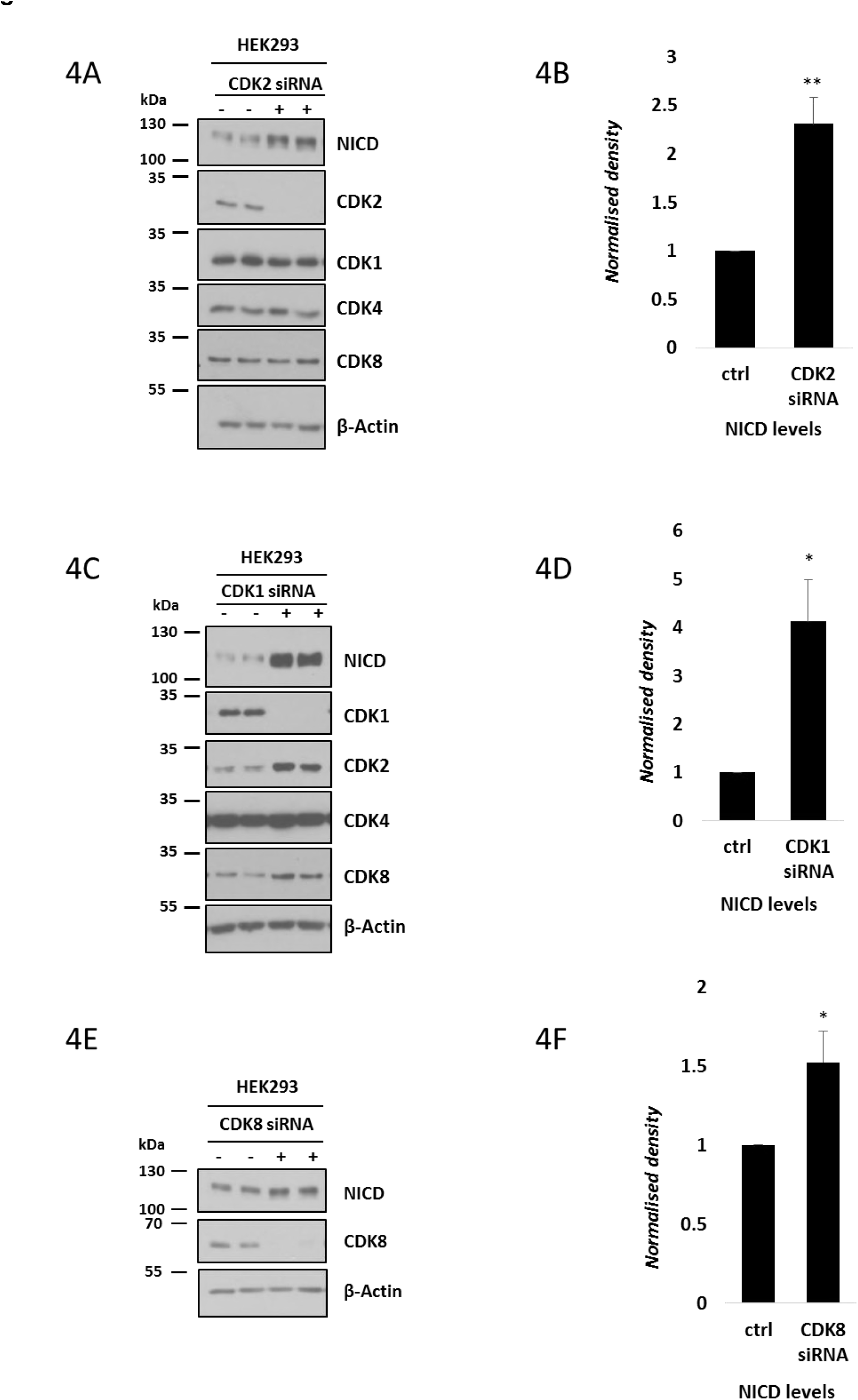
CDK1, CDK2 and CDK8 depletion increased endogenous levels of NICD in HEK293 cells. (A) HEK293 cells were cultured for 48h after transfection with plasmids encoding scrambled siRNA (-) or siRNA specific for CDK2 (+) followed by western blot for NICD, CDK2, CDK1, CDK4 and CDK8. β-Actin served as loading control. (B) Quantification of the density of western blot bands in (A) performed by ImageJ software. Data are expressed as fold changes compared to control (scrambled siRNA transfected) cell lysate. All data represent the mean ± SEM from three independent experiments. Student’s t-test analysis was performed, with **p≤0.01. (C) HEK293 cells were cultured for 48h after transfection with plasmids encoding scrambled siRNA (-) or siRNA specific for CDK1 (+) followed by Western blot for NICD, CDK1, CDK2, CDK4 and CDK8. β-Actin has been used as loading control. (D) Quantification of the density of western blot bands in (C) performed by ImageJ software. Data are expressed as fold changes compared to control (scrambled siRNA transfected) cell lysate. All data represent the mean ± SEM from three independent experiments. Student’s t-test analysis was performed, with *p≤0.05. (E) HEK293 cells were cultured for 48h after transfection with plasmids encoding scrambled siRNA or siRNA specific for CDK8, followed by western blot for NICD and CDK8. β-Actin has been used as loading control. (F) Quantification of the density of western blot bands in (E) performed by ImageJ software. Data are expressed as fold changes compared to control (scrambled siRNA transfected) cell lysate. All data represent the mean ± SEM from three independent experiments. Student’s t-test analysis was performed, with *p≤0.05.

We repeated this assay and depleted CDK1 by a siRNA-mediated approach in HEK293 cells. Under these conditions we also observed an increase in NICD protein levels compared to the control (**Figure 4C**), and this increase was statistically significant (**Figure 4D**).Interestingly, we also detected elevated levels of CDK2 and CDK8 following CDK1 depletion, suggesting a possible compensation effect was occurring upon CDK1 knockdown. Nevertheless, this did not prevent the effect of loss of CDK1 upon NICD turnover.

CDK8 has previously been proposed as a potential kinase involved in NICD phosphorylation and turnover [35, 65]. We monitored NICD levels following efficient siRNA-mediated depletion of CDK8 in HEK293 cells. This resulted in very slightly elevated NICD levels (**Figures 4E-F**). These data suggest CDK8 may also phosphorylate NICD and regulate its turnover as previously proposed [35, 65].

Taken together, these data provide further validation of CDK1 and CDK2 involvement in NICD phosphorylation and turnover.

### Pharmacological inhibition of CDK2 and CDK1 activity increases levels of NICD *in vitro*

As a complementary approach to siRNA-mediated loss of function, we selected two small molecule inhibitors that have a highly selective inhibitory activity against CDK2 kinase: (Purvalanol B and GSK650394A). The specificity of each of these kinase inhibitors has been tested (at two different concentrations) by the International Centre for Kinase profiling within the MRC Protein Phosphorylation Unit at the University of Dundee. At 1 and 10 μM, Roscovitine specifically inhibits more than 96% of CDK2 activity. Purvalanol B inhibits more than 95% CDK2 activity, at 0.1 and 1 μM. GSK650394A inhibits 99% of CDK2 activity at both 1 and 10 μM (**Figure 5A**). CDK2 is by far the most sensitive target for all three inhibitors (**Figure 5A**). HEK293 cells treated with Roscovitine, Purvalanol B or GSK650394A (10 μM, 0.1 μM and 10 μM, respectively) for 3 hours exhibited significantly increased NICD levels compared to control DMSO treated cells (**Figures 5B-C**). Under these conditions protein levels of both CDK2 and Cyclin E were not affected (data not shown). iPS cells cultured 3 hours in the presence of 0.1 μM Purvalanol B also exhibited significantly elevated levels of NICD as compared to DMSO treated control cells, indicating this is a conserved effect of CDK2 (**Supplementary Figures 4A-B**).

**Figure 5.**
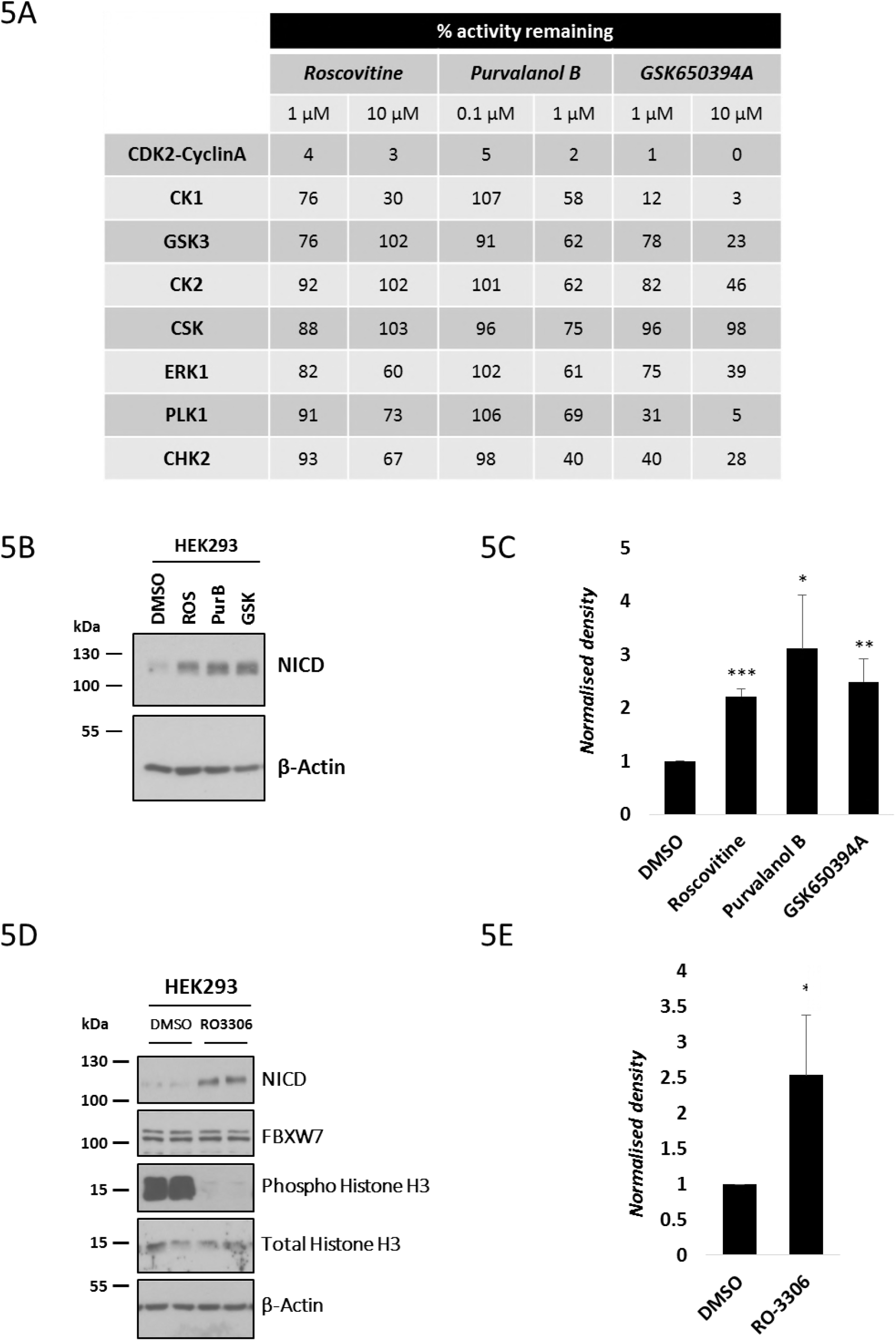
Pharmacological inhibition of CDK1 and CDK2 leads to increased NICD levels *in vitro*. (A) Analysis of the inhibitory activity of three highly selective CDK2 inhibitors against a panel of kinases - a selection of those tested is shown here. At both 1 and 10μM, Roscovitine is able to inhibit more than 95% of CDK2 activity, but is far less effective against other kinases. Purvalanol B (0.1 and 1μM) inhibits more than 94% of CDK2 activity. At both 1 and 10 μM, GSK650394A is able to inhibit more than 98% of CDK2 activity. Source: The Kinase Profiling Inhibitor Database (http://www.kinase-screen.mrc.ac.uk/kinase-inhibitors). (B) HEK293 cells were treated with three highly selective CDK2 inhibitors (10 μM of Roscovitine, 0.1μM of Purvalanol B and 10μM of GSK650394A) for 3 hours. Endogenous levels of NICD were detected by western blot. β-Actin served as loading control. (C) Quantification of the density of western blot bands in (B) using ImageJ software. Data are expressed as fold changes compared to DMSO. All data represent the mean ± SEM from three independent experiments. Student’s t-test analysis was performed, with *p≤0.05, **p≤0.01, and ***p≤0.001. (D) HEK293 cells were treated with RO-3306, a specific CDK1 inhibitor, for 3 hours at 10 μM. Levels of NICD, FBXW7, phospho-Histone H3 and Histone H3 were detected by western blot. β-Actin was used as loading control. (E) Quantification of the density of western blot bands in (D) using ImageJ software. Data are expressed as fold changes compared to DMSO. All data represent the mean ± SEM from three independent experiments. Student’s t-test analysis was performed, with *p≤0.05.

Given our observation that both CDK1 and CDK2 can phosphorylate NICD peptides *in vitro*, we similarly treated HEK293 cells with a CDK1 specific inhibitor, RO-3306 [68]. Following exposure to 10 μM of RO-3306 for 3 hours HEK293 cells showed elevated NICD levels compared to control DMSO treated cells (**Figures 5D-E**). These data further support the hypothesis that both CDK1 and CDK2 are likely to be involved in phosphorylation-mediated regulation of NICD turnover.

### NICD levels fluctuate over the cell cycle

It is well known that CDK1 and CDK2 share several substrates, with a consequent functional redundancy [69]. Our data demonstrate that these two kinases can phosphorylate NICD *in vitro* and in the absence of either kinase NICD levels increase in HEK293 cells suggesting they are not acting redundantly in this context. In order to further test this, we analysed whether NICD levels fluctuate during the cell cycle where the role of both CDK1 and CDK2 has been extensively reported in regulating transition to distinct cell cycle phases [70].

To that end, we synchronised HEK293 cells by using a double thymidine block assay. After releasing from the second thymidine block, cells were collected at indicated time points and cell cycle characterization was performed by Fluorescence Activated Cell Sorting (FACS) (**Figure 6A**). **Figure 6A** shows the distribution though the cell cycle of HEK293 cells after synchronization at early S-Phase (0 hour), as previously reported [71]. Two and four hours post release, the majority of cells were in S-phase. At six hours post release, the majority of cells were in G2 phase, while eight hours after release, the majority of cells were in late G2/early M phase. At ten hours post release, the majority of cells had exited mitotic phase and already entered G1. After twelve hours from release, cells were in mid/late G1 of the new cell cycle (**Figure 6A**). The graph in **Figure 6B** represents the cell cycle distribution of HEK293 cells at distinct time points post double thymidine block and release from three independent experiments analysed by FACS. Western blotting analysis of synchronized HEK293 cell extracts showed the expression of several cell cycle regulatory proteins at the same distinct time points reflecting distinct cell cycle phases as described above (**Figure 6C**). Interestingly, we found that NICD levels fluctuated in a striking manner during the cell cycle, whereas we saw no change to levels of FBXW7. At 2, 4, 8 and 12 hours post release, we observed a dramatic decrease of NICD expression corresponding to CDK2-dependent G1/S phase, whereas the decrease 8 hours after double thymidine block release occurred in the CDK1-dependent G2/M phase transition. These data suggest that NICD levels fluctuate during the cell cycle in a CDK1 and CDK2-dependent manner.

**Figure 6.**
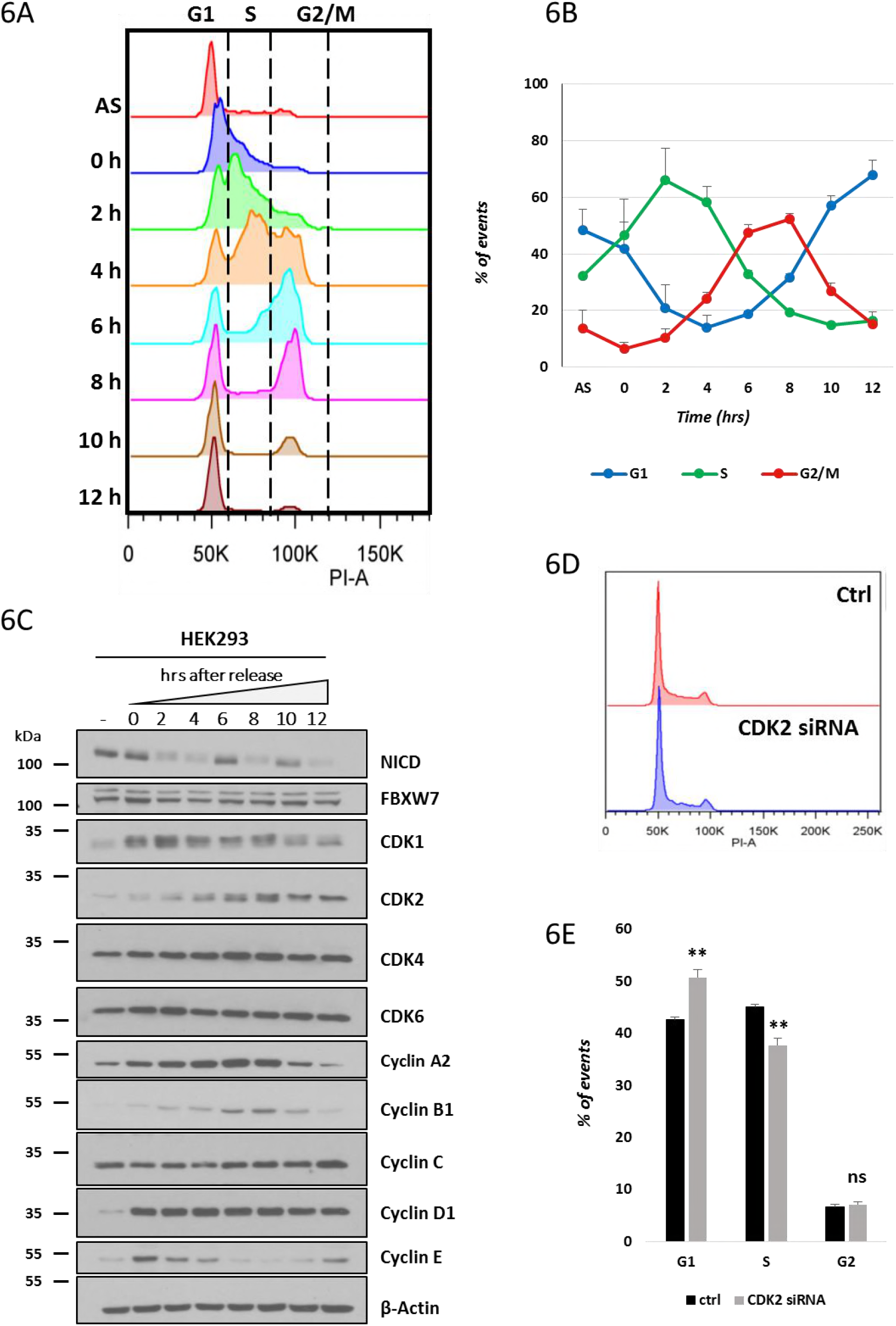
NICD levels fluctuate during the cell cycle. (A) Cell cycle profile for HEK293 cells released from synchronization after double thymidine block. Cells were released and harvested at the indicated time points (AS= asynchronous). Analysis of cell cycle arrest and release was performed using propidium iodide (PI) staining and flow cytometry. A representative experiment of three performed is shown. (B) Chart of flow cytometry data shows the percentage of HEK293 cells in G1, S and the G2/M phases after release from double thymidine treatment. Time points are expressed as mean ± SEM from three independent experiments. (C) Expression of the indicated proteins in HEK293 cells was examined by western blotting, and β-Actin was used as loading control. This summary is a representation of three independent experiments. (D) Cell cycle profile for HEK293 cells 48h after transfection with plasmids encoding scrambled siRNA or siRNA specific for CDK2. Analysis of cell cycle arrest and release was performed using propidium iodide (PI) staining and flow cytometry. A representative experiment of three performed is shown. (E) Graph of flow cytometry data shows the percentage of cells in given cell-cycle phases 48h after transfection with plasmids encoding scrambled siRNA or siRNA specific for CDK2. Graph represents the mean of three independent experiments. All data represent the mean ± SEM from three independent experiments. Student’s t-test analysis was performed, with **p≤0.01 (ns=not significant).

In addition, we analysed the cell cycle distribution by FACS in HEK293 cells after CDK2-siRNA mediated depletion, which we have shown leads to a significant increase in NICD levels (**Figure 4A**) and we observed a statistically significant accumulation of cells in G1 phase compared to the control, as expected for cells deprived of CDK2 activity and therefore unable to pass the G1/S checkpoint (**Figures 6D-E**). These data suggest that the drop in NICD levels occurring in G1/S phase is due to CDK2 phosphorylation of NICD.

### Pharmacological inhibition of CDK2 increases NICD levels and delays the pace of the segmentation clock in mouse PSM explants

In order to address whether CDK2 phosphorylation of NICD is involved in driving the NICD-FBXW7 interaction, we performed a co-immunoprecipitation assay with FBXW7 antibody and analysed NICD by western blot after CDK2 inhibitor treatment. As above, in order to maximise the amount of NICD immunoprecipitated HEK293 cells were treated with MLN4924 (to prevent NICD degradation) +/- Purvalanol B (0.1 μM). A significantly reduced interaction between NICD and FBXW7 was observed after treating HEK293 cells with Purvalanol B for 3 hours. This did not reflect a reduction in the level of immunoprecipitated FBXW7 (**Figure 7A**). Statistical analysis on the density of western blot bands after immunoprecipitation, confirmed an extremely significant reduction in the NICD-FBXW7 interaction following Purvalanol B treatment (**Figure 7B**).

**Figure 7.**
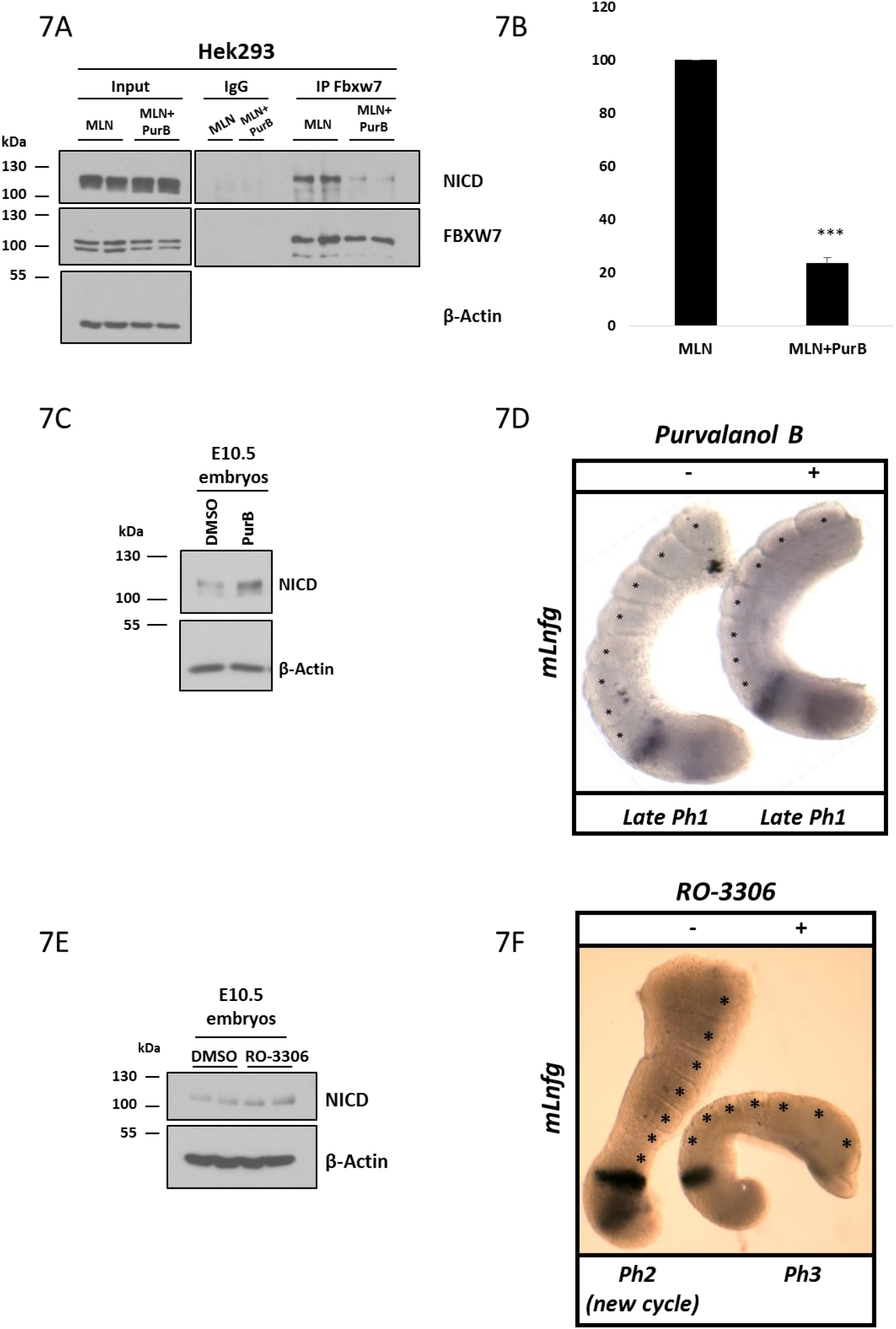
Purvalanol B treatment reduces NICD-FBXW7 interaction and delays the pace of the segmentation clock in mouse PSM explants. (A) Purvalanol B treatment reduced the NICD-FBXW7 interaction. 200 μg of HEK293 cell lysates treated with MLN4924 or MLN4924 in combination with 0.1 μM of Purvalanol B were subjected to immunoprecipitation using FBXW7 antibody, or IgG antibody as negative control, and precipitated material was analysed by western blot using NICD antibody. Western blot with FBXW7 antibody served as loading control for immunoprecipitation efficiency. 10% of cell lysate before immunoprecipitation was used as input control and β-Actin has been used as loading control. (B) Quantification of the density of western blot bands in (A) performed by ImageJ software. Data are expressed as fold changes compared to MLN4924 treated samples. All data represent the mean ± SEM from three independent experiments. Student’s t-test analysis was performed, with ***p≤0.001. (C) E10.5 mouse tails were bissected down the midline. One half (+) was cultured for 4 hrs in the presence of Purvalanol B (1 μM). The contralateral half (-) was cultured for 4h in the presence of DMSO. Control or treated explants were pooled and NICD levels were detected by western blot. β-Actin was used as loading control. (D) Bissected E10.5 mouse PSM explants were cultured in the absence (-) or presence (+) of 1μM of Purvalanol B for 4 hours and then analysed by *in situ* hybridization for *mLfng* mRNA expression. Purvalanol B treated explant has one less somite than the control explant and the treated explant is in the same late phase 1 of the oscillation cycle of dynamic *mLfng* mRNA expression indicating it is a whole cycle delayed compared to the “-” explant. (E) E10.5 mouse tails were bissected down the midline. One half (+) was cultured for 4 hrs in the presence of RO-3306 (10 μM). The contralateral half (-) was cultured for 4h in the presence of DMSO. Control or treated explants were pooled and NICD levels were detected by western blot. β-Actin has been used as loading control. (F) Bissected E10.5 mouse PSM explants were cultured in the absence (-) or presence (+) of 10 μM of RO-3306 for 4 hours and then analysed by *in situ* hybridization for *mLfng* mRNA expression. RO-3306 treated explant has one less somite than the control explant and the treated explant is in a phase behind of the oscillation cycle of dynamic *mLfng* mRNA expression indicating there is a delay in the oscillation compared to the “-” explant.

In order to address the potential *in vivo* role of CDK-mediated NICD phosphorylation during somitogenesis, we cultured E10.5 mouse PSM explants for 4 hours in the presence of 1μM of Purvalanol B. Initially we analysed NICD levels by western Blot and just as in the *in vitro* context, CDK2 inhibition resulted in increased NICD levels as compared to control embryos (**Figure 7C**). This provides the first *in vivo* evidence CDK2 is likely to be involved in NICD turnover.

Previous reports have suggested perturbations to NICD turnover leading to increased NICD levels/stability are closely linked to an increase in the period of segmentation clock oscillations in the PSM [18]. To further explore whether clock gene oscillations were delayed following CDK2 inhibition, we used the half-embryo assay, where the PSM from one half of an E10.5 mouse embryo is cultured in control media, while the contralateral half from the same embryo is cultured in the presence of 1μM of Purvalanol B for 4 hours. In 55.5% of cases examined (n=10/18), exposure of PSM explants to Purvalanol B caused a delay in the pace of oscillatory *mLfng* expression across the PSM as compared to the control explant (**Figure 7D**). Also, in some cases the treated explant (“+”) developed one somite less compared to the control (**Figure 7D**).

Additionally, we cultured E10.5 mouse PSM explants in the presence or absence of the CDK1 inhibitor, RO-3306, for 4 hours at 10μM. Analysis of NICD levels by western blot revealed that inhibition of CDK1 leads to elevated levels of NICD compared to control embryos (**Figure 7E**). E10.5 mouse half embryo explants exposed to RO-3306 treatment for 4 hours showed delayed clock oscillations of *mLfng* as compared to DMSO treated contralateral half embryo explants (**Figure 7F**). These data, not only support our *in vitro* findings, but importantly provide, for the first time, the *in vivo* evidence that CDK1 and CDK2 are involved in NICD stability and turnover and that this molecular regulation of NICD turnover is extricably linked to the pace of the segmentation clock.

### Mathematical model links NICD regulation and cell cycle

To understand how our findings on the molecular details of NICD regulation in individual cells give rise to tissue-scale delay of the segmentation clock, we first developed a mathematical model of NICD production and degradation in HEK293 cells. The variables in the model define the position of a cell in the cell cycle, the amount of NICD and the amount of phosphorylated NICD (pNICD) at time *t*. The model enables us to connect assumptions about molecular processes in individual cells with experiments performed on a large population of cells.

We assume a *K*-compartment model that describes sequential progression of a cell through the cell cycle (**Figure 8A**). It is assumed that CDKs that phosphorylate NICD, resulting in its interaction with FBXW7 and subsequent degradation, are active in some *m* of the cell cycle states. We assume that: (i) NICD exists in non-phosphorylated and phosphorylated forms; (ii) NICD is produced at constant rate *k*_1_; (iii) both forms of NICD get degraded at background rate *k*_7_; (iv) NICD gets phosphorylated by CDKs at rate *k*_3_; (v) pNICD degrades at rate *k*_6_ with *k*_6_ > *k*_7_; and (vi) dephosphorylation of NICD occurs at rate *k*_2_. We consider the pharmacological perturbations as follows: LY411575 treatment experiments, in which NICD production is inhibited, are simulated by setting *k*_1_ = 0; Purvalanol B/Roscovitine treatment experiments, which target only CDK2, reduce the number of compartments where CDKs are active by a factor of two; and MLN4924 treatment, which inhibits FBXW7 mediated degradation of NICD, are simulated by removing the fast mode of decay (*k_6_* = 0).

**Figure 8.**
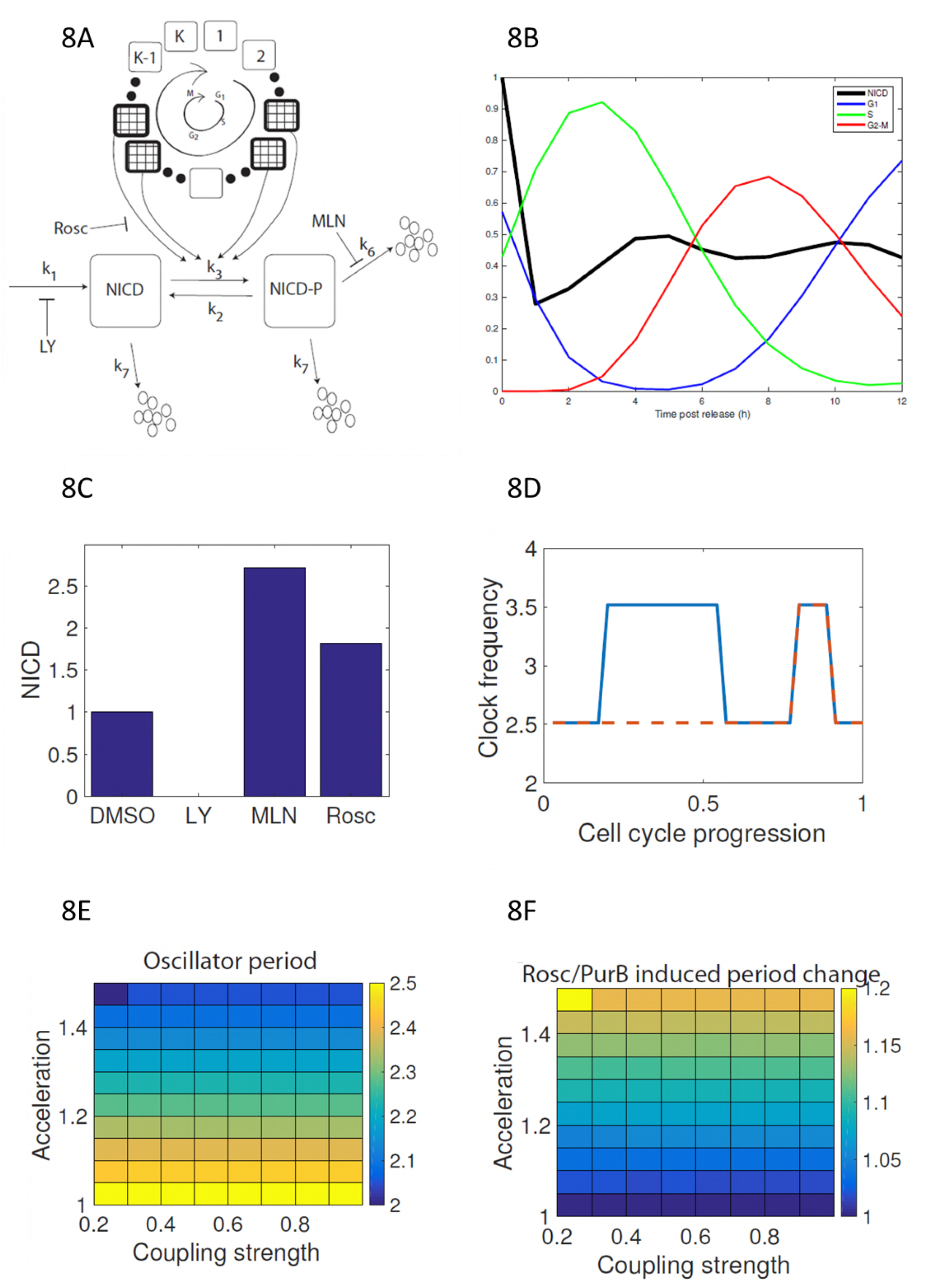
Mathematical model links NICD regulation, segmentation clock and cell cycle. (A) A schematic illustration of NICD production, phosphorylation and cell-cycle coupled degradation. The cell cycle is represented by clockwise progression through a *K* state model. NICD phosphorylation occurs at two different stages of the cell cycle (patterned boxes). (B) Levels of NICD and cell cycle phase distribution are plotted against time post thymidine release. (C) Steady-state levels of NICD are plotted for simulated control, LY411575, MLN4924 and Purvalanol B/Roscovitine treatments. (D, E, F) Simulating a population of coupled PSM oscillators with cell cycle modulated frequencies. (D) The clock frequency in a single cell is plotted against cell cycle position for normal (solid line) and CDK2 inhibited (dashed line) cells. (E) The emergent population-scale oscillator period is plotted against coupling strength and CDK acceleration factor. (F) Period change (Roscovitine/Purvalanol B versus control) as a result of CDK inhibition is plotted against coupling strength and CDK-mediated acceleration rate.

By averaging cellular descriptions of NICD and the cell cycle over a given cell population, we derive equations describing the average levels of total NICD (i.e. as measured in a Western Blot experiment, see Methods). Upon release from double thymidine block, cells are synchronised in the cell cycle and there are relatively high levels of CDK activity at different times post release. As pNICD degrades faster, high levels of CDK results in lower levels of NICD (see **Figure 8** and **Figures 4A-D,5B-E, 6C, 7C-E**). We find the model qualitatively fits the experimental observations if NICD is phosphorylated in two separate time windows post release. We propose that these windows correspond to activity of CDK2 (G1/S phase) and CDK1 (M phase) (**Figures 8A and 8B**). In a situation where the cell cycle is desynchronised, the population-averaged phosphorylation rate (*mk*_3_/*K*, where *m*/*K* is the fraction of the cell cycle where CDKs are active) is a fraction of the true phosphorylation rate. The model can reproduce Western Blot results from DMSO and MLN4924 experiments (**Figure 8C**), allowing inference of the parameters *k*_2_ and *k*_3_. Notably, the simulated Purvalanol B/Roscovitine treatment experiment presented in Figure 8C is a prediction that is validated by the experimental data.

To explore how CDK inhibition results in delay of the segmentation clock we introduce an additional variable representing phase of the somitogenesis clock and assume that the caudal PSM behaves like a population of phase coupled oscillators. In Wiederman et al. we proposed a model that showed how competition between activators and inhibitors of clock gene transcription could lead to the oscillator period being an increasing function of NICD half-life [18]. Here, we impose, without explicitly describing the molecular circuitry, this assumption. We assume that each cellular oscillator has a natural frequency *ω_i_(t)* that is a function of cell cycle position; when CDKs are active the baseline clock frequency, *ω_0_*, is increased by an acceleration factor *r* (**Figure 8D**). As the cell cycle is asynchronous in the PSM, it is therefore comprised of a population of oscillators with frequencies *ω*_0_ or *ωr*_0_. Given sufficiently strong coupling a population of such oscillators yields synchronous oscillations with a period that is an average of the individual oscillator periods (**Figures 8E-8F**). Upon CDK inhibition, the relative number of faster oscillators is reduced hence the average period decreases (**Figure 8E**). Hence the cell cycle somitogenesis coupled model provide a mechanistic description of how CDK-mediated phosphorylation of NICD can result in the observed phenotype in PSM tissue (**Figure 7**).

## DISCUSSION

The Notch pathway plays multiple critical roles in somitogenesis. Notch signalling is critical for dynamic clock gene expression and for somite formation in mice [9]. Notch-Delta signalling is essential to synchronise clock gene oscillations among neighbouring cells of the PSM [10–13]. Moreover, a recent publication using a pharmacological approach, has shown that modulating the half-life of NICD affects the clock oscillation period and somite size [18]. It is important to note that NICD degradation and turnover still occurred under these conditions but less efficiently. In this study, we have significantly extended those findings to determine the mechanistic details of regulation of NICD turnover, and we report for the first time that cell cycle-dependent CDK1 and CDK2 activity is involved in NICD turnover, which has broad implications across all developmental and disease contexts where Notch plays a role.

We demonstrate that kinase inhibitors (Roscovitine, DRB and XAV939), previously shown to prolong NICD half-life and delay the segmentation clock pace in mouse and chick PSM *in vivo*, [18] also increased endogenous levels of human NICD when a range of primary human cell lines were treated for 3 hours. This highlights the conserved effect of these inhibitors on regulating NICD stability, which is perhaps not surprising given the high degree of sequence similarity between mouse, human and chicken NICD.

We demonstrate that, following exposure to these kinase inhibitors, the increased stability of NICD accompanies a change in the array of phosphorylated isoforms of NICD observed. The different profiles of phosphorylated isoforms of NICD observed, following exposure to each of the inhibitors, suggest they each inhibit distinct kinases. In each case, however, the highest molecular bands were no longer visible. It is possible that the multiple phosphorylation bands are indicative of unique events, some of which may reflect priming phosphorylation events that facilitate or increase the efficiency of secondary phosphorylation events, which then act as phospho-degron signals to recruit E3 ligases that target NICD for degradation. It is not possible to determine whether the lower molecular weight bands that remain, following inhibitor treatment, are indicative of the loss of primary or secondary phosphorylation events.

A number of reports have highlighted the fact that the SCF^FBXW7^ E3 ligase plays an important role in NICD degradation [35, 44, 46–48, 66]. SCFs (Skp1, Cullin-1, F-box protein) are a class of E3 ligases that use Cullin-1 as a scaffold and F-box proteins as substrate receptors. FBXW7 is an evolutionary conserved F-box protein. Substrate phosphorylation instigates FBXW7 binding to a conserved CDC4 phospho-degron motif which then recruits the rest of the E3 ligase complex, including Cullin1, thereby targeting the substrate for ubiquitination and subsequent degradation by the proteasome [45, 49]. The role of FBXW7 in NICD regulation is also supported by data showing that, when mammalian Sel-10 (homologue of FBXW7 in *C. elegans*) is mutated, NICD is much more stable [46]. To date, the interaction between NICD and FBXW7 by co-immunoprecipitation has only been shown using overexpressed proteins, due to the fact this interaction is very transient and leads to efficient degradation of NICD [35, 44, 47, 48, 66]. In order to stabilise the interacting complex, we conducted experiments in the presence of MLN4924, a Cullin1 neddylation inhibitor which prevents activation of Cullin1. Under these conditions, we demonstrate for the first time that NICD and FBXW7 can interact at endogenous levels in HEK293 cells. Moreover, this allowed us to demonstrate that, when phosphorylation is disrupted by small molecule CDK inhibitors (Roscovitine or DRB), the NICD-FBXW7 interaction is reduced. It is important to note that neither inhibitor abolished the NICD-FBXW7 interaction which suggests again that they are each inhibiting only some of the kinase activity involved in NICD phosphorylation and subsequent recruitment of FBXW7. This aligns with the observation that in the chick/mouse PSM both of these inhibitors increase NICD stability and the period of the segmentation clock but that NICD turnover still occurs in this tissue, and thus is dependent on a number of different kinases/phosphorylation events differentially targeted by these two inhibitors [18].

By Mass Spectrometry analysis, we identified 15 phospho-sites within human exogenous NICD in HEK293 cells, some of which have been previously identified and reported to be involved with NICD turnover [35, 65]. Among those phospho-sites identified, we found that Serine 2513, when mutated to alanine, thereby rendering the site non phosphorylatable, was essential for the interaction between NICD and FBXW7. Point mutations in a number of other phosphorylated residues showed these are non-essential for this NICD-FBXW7 interaction, including two residues that have previously been reported to potentially be required for the interaction; S2484 and S2506 (S2516 and S2538 in our annotation) [35, 66]. However, Fryer and colleagues used exogenous proteins and mutated all three sites simultaneously so the individual contribution/requirement of each residue was not addressed in that study. O’Neil and colleagues have also reported, using exogenous proteins, that phosphorylation of the Threonine residue at T2487 in mouse NICD (T2511 in our annotation) is key to driving the NICD-FBXW7 interaction and subsequent ubiquitination of NICD [44]. We did not observe phosphorylation at this residue in human NICD. However it would be interesting to repeat the mass spec analysis using different enzyme digestions to determine if this reveals additional phosphorylated sites that play a role in this interaction.

SCF substrates typically have many phospho-degrons that can be widely dispersed. It will be important to test the requirement of all the phosphorylated residues we have identified in NICD for driving and regulating the NICD-FBXW7 interaction.

We have identified CDK1 and CDK2 as two kinases that can phosphorylate NICD in the PEST region that harbours residue serine 2513, which we have demonstrated to be crucial for the interaction between NICD and FBXW7. Through two loss of function approaches, namely siRNA and a pharmacological approach, we further demonstrated the role of both CDK1 and CDK2 kinases in the regulation of NICD turnover. Indeed, by transfecting HEK293 cells with CDK1, or CDK2 specific siRNA, or treating HEK293 cells or iPS cells with highly selective small molecule inhibitors against CDK1 and CDK2, we could appreciate an increase in NICD levels, indicating that inhibiting phosphorylation by CDK1 or CDK2, renders NICD more stable. As described above, the serine 2513 residue has previously been reported to potentially be involved in NICD turnover in other systems [35]. Using gain and loss of function experiments with exogenous proteins, the authors reported phosphorylation occurred through CyclinC:CDK8. Indeed, by transfecting HEK293 cells with CDK8 specific siRNA, we could also appreciate an increase in NICD levels, indicating that this kinase could also play a role in NICD turnover, although the effect appears less dramatic compared to that observed with CDK1 or CDK2 specific siRNA.

CDK1 and CDK2 activity and thus phosphorylation of their substrates changes in a cell cycle dependent manner [72]. Strikingly, we find that NICD levels vary in HEK293 cells in a cell cycle-dependent way such that we observe lowest levels of NICD in phases of the cell cycle where CDK1/Cyclin B1 and CDK2/Cyclin E levels are highest and therefore when these complexes are reported to be most active [73]. This finding suggests that NICD activity, signal duration and signal strength is likely to vary in a cell cycle dependent manner, potentially leading to differential transcriptional output in different phases of the cell cycle.

Given our findings, it is striking that CDK2 homozygous *null* mice are viable [74]. However, it has been reported that CDK1 and CDK2 share more than 50% of their targets [75] which is likely to allow for a lot of redundancy in this knock-out line, as CDK1 can compensate for loss of CDK2 by forming active complexes with A-, B-, E- and D-type cyclins. The CDK1-*null* homozygous mice however are embryonic lethal and a CDK2 knock-in to the CDK1 locus is unable to rescue this phenotype [76, 77]. It would be interesting to examine the role of the CDK1 conditional mutant in the context of Notch signalling and somitogenesis [78]. Interestingly, Cyclin C-*null* mice, which are embryonic lethal at 10.5, show dramatically reduced NICD1 phosphorylation *in vivo*, and elevated NICD1 levels. The authors show that Cyclin C can complex with CDK3, CDK19, CDK8, CDK1 and CDK2 to phosphorylate NICD1 and promote NICD1 degradation [65]. Thus, it would be very interesting to examine a PSM conditional Cyclin C loss of function to determine whether this cyclin is involved in regulating NICD turnover in this tissue. It is noteworthy Cyclin C levels did not appear to vary in a cell cycle dependent manner in HEK293 cells, although it is possible the activity of Cyclin C/CDK complexes varies in a cell cycle-dependent manner through post-translational modifications rather than protein levels per se.

The finding that CDK1 inhibition or CDK2 inhibition leads to increased levels of NICD in E10.5 mouse embryo PSM lysates and to a delay in clock gene oscillations and somite formation provides the first *in vivo* evidence that both CDK1 and CDK2 are involved in NICD turnover.

It is noteworthy that inhibition of CDK2 activity with a highly selective inhibitor reduces the NICD-FBXW7 interaction in HEK293 cells but does not block it completely, as we saw with Roscovitine and DRB. Indeed, inhibition of CDK1 or CDK2 in the mouse PSM with highly selective inhibitors raises NICD levels and also slows, but does not block, dynamic Notch target clock gene expression, and somitogenesis. This suggests that some NICD turnover persists under these conditions, possibly through redundancy between CDK1 and 2 and/or through CDK8-mediated phosphorylation, the subsequent recruitment of E3 ligases and degradation of NICD. Moreover, a report by Chiang *et al*. has also identified a region downstream of the PEST sequence, termed S4, that is involved in NICD degradation, but that is independent of FBXW7 activity [79]. These data indicate there are several mechanisms regulating NICD turnover which are partially redundant.

We also developed a mathematical model that coupled cell cycle dynamics to NICD degradation. Using HEK293 cells, model parameters were identified that recapitulated the distributions of cells in the different cell cycle phases. To recover qualitative features of NICD thymidine release experiments, NICD phosphorylation was required at distinct stages in the cell cycle. After using MLN4924 and control experiments to estimate NICD phosphorylation rate the model qualitatively predicts the effect of Roscovitine/PurB treatment on total NICD levels. To address the likely effect of CDK2 inhibition in the PSM, the model was extended to account for position of a cell in the segmentation clock cycle. Following Wiederman *et al*. [18], it was assumed that levels of NICD are anti-correlated with clock somitogenesis frequency. Hence, the posterior PSM is represented by a population of phase coupled oscillators whose frequency is cell cycle dependent. Simulating CDK2 inhibition removes a pool of faster oscillators thus reducing the tissue period.

Notch plays a key role as a gatekeeper protecting progenitor and/or stem cells in multiple developmental contexts, in part through preventing differentiation and in part through regulating components of the cell cycle [80–82]. Our novel finding of a reciprocal auto-regulatory role between the cell cycle regulated CDKs, CDK1 and CDK2, and NICD turnover has potentially great relevance to the developmental biology community and may provide additional insight into disease/cancer contexts where this autoregulation may have gone awry.

## MATERIALS AND METHODS

All plasmids and reagents indicated as such in the text are available from MRC PPU Reagents and Services (https://mrcppureagents.dundee.ac.uk/)

### Cell Culture

HEK293 (human embryo kidney cells) and IMR90 (Human Caucasian fetal lung fibroblast) cells were obtained from the American Type Culture Collection (ATCC). Cells were routinely cultured and maintained in DMEM High glucose (Gibco) supplemented with 10% Fetal Bovine Serum (FBS; LabTech), 1% Penicillin/Streptomycin (Gibco), 1% Sodium Pyruvate (Gibco) and 2mM L-Glutamine (Gibco).

ChiPS4 human Induced pluripotent stem (iPS) cells (derived from new born human dermal fibroblasts) were purchased from Cellartis AB and maintained using DEF-CS (Cellartis AB) according to the manufacturer’s recommendations. For experiments, cells were seeded as single cells on Geltrex^TM^ coated dishes (10 μg/cm^2^) in DEF medium supplemented with 10 μM of Rho-kinase inhibitor Y27632 (Tocris) at a density of 1×54 cells/cm^2^ and allowed to attach overnight. The medium was then replaced with fresh DEF medium (Y27632) and after a further 24 hrs cells were treated with inhibitors.

All cells were grown at 37°C in 5% CO_2_.

Cells were trypsinized and split into new plates at subconfluency.

### Transfections

#### Overexpression transfections

HEK293 cells were seeded 24 hrs prior to transfection and then transiently transfected following the GeneJuice^®^ Transfection Reagent protocol provided by Merck Millipore. Depending on plate size, 2-9 μg of plasmid was used. Cells were harvested 24 hrs after transfection.

### siRNA transfections

Small interfering RNA oligonucleotides were purchased from MWG/Eurofins and used in a stock concentration of 20μM. siRNAs were transfected using calcium phosphate transfection method [83]. siRNA oligonucleotides were diluted in a solution containing distilled sterile H_2_O and 2M CaCl_2_.

The mixture was then added drop-wise to a round-bottom 15ml tube containing 2X HBS (0.137M NaCl, 0.75M Na2HPO4, 20mM HEPES pH 7.0). The content was then added to cells and incubated overnight at 37°C. Media was changed after 24 hrs and cells were harvested after 48 hrs of transfection.

### Oligonucleotide Sequences for siRNA Knockdown siRNA siRNA

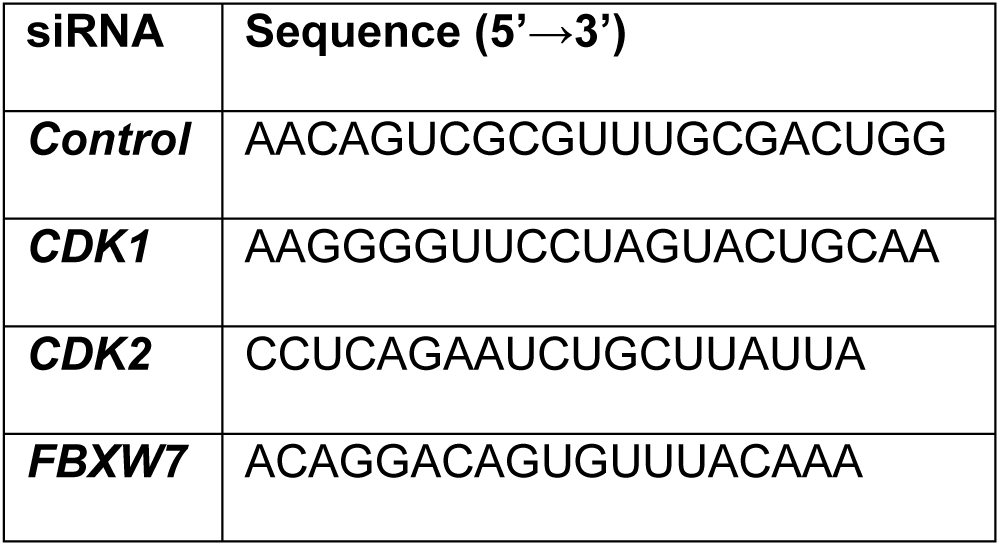

Commercial CDK8 siRNA (Cell Signalling, #6438) has been used.

### Plasmids and mutagenesis

hNICD-GFP vector was generated and obtained from MRC-PPU reagents, University of Dundee. All NOTCH1 mutants were generated and obtained from MRC-PPU reagents, University of Dundee.

Briefly, the fragment NOTCH1 1754-2555 (end) was synthesized by GeneArt with flanking BamHI and NotI restriction sites to facilitate cloning, the sequence was codon-optimized for mammalian expression. This was then digested and ligated into expression vector pCMV5D GFP to make the wild type clone pCMV5D GFP NOTCH1 1754-end.

Site-directed mutagenesis was carried out using the QuikChange Lightning Site-Directed Mutagenesis Kit (Agilent Technologies) but substituting the Taq with KOD Hot Start DNA polymerase (Novagen). All mutations were confirmed by sequencing.

S2205A change was introduced using primers

Forward: 5’-CCCGTGGATAGCCTGGAAGCGCCTCACGGCTACCTGAGC

Reverse: 5’-GCTCAGGTAGCCGTGAGGCGCTTCCAGGCTATCCACGGG

S2513A change was introduced using primers:

Forward: 5’-CCCATTTCTGACCCCTGCACCCGAGAGCCCCGATC

Reverse: 5’-ATCGGGGCTCTCGGGTGCAGGGGTCAGAAATGGG

S2516A change was introduced using primers

Forward: 5’-CTGACCCCTAGCCCCGAGGCGCCCGATCAGTGGTCTAGC

Reverse: 5’-GCTAGACCACTGATCGGGCGCCTCGGGGCTAGGGGTCAG

S2513A and S2516A double mutant was generated with primers:

Forward: 5’-CACCCCTTCCTCACCCCGGCCCCTGAGGCCCCTGACCAGTGGTCCA

Reverse: 5’-TGGACCACTGGTCAGGGGCCTCAGGGGCCGGGGTGAGGAAGGGGTG

S2527A change was introduced using primers

Forward: 5’-TCTAGCAGCAGCCCCCACGCGAACGTGTCCGATTGGAGC

Reverse: 5’-GCTCCAATCGGACACGTTCGCGTGGGGGCTGCTGCTAGA

S2538A change was introduced using primers

Forward: 5’-TGGAGCGAAGGCGTGTCCGCCCCCCCAACCAGCATGCAG

Reverse: 5’-CTGCATGCTGGTTGGGGGGGCGGACACGCCTTCGCTCCA

### Treatments

Cells were treated for 3 hrs with different drugs: 150 nM of LY411575 (generated in house, University of Dundee) [9], 1μM of MLN4924 (MRC-PPU reagents, University of Dundee), 10 μM of 5,6-Dichloro-1-beta-D-ribofuranosylbenzimidazole (DRB) (Sigma), 10 μM of Roscovitine (Calbiochem), 10 μM of XAV939 (Tocris Bioscience), 0.1 μM of Purvalanol B (MRC-PPU reagents, University of Dundee), 10 μM of GSK650394A (MRC-PPU reagents, University of Dundee), 10 μM of RO-3306 (Sigma) and DMSO (Sigma) as control.

Lysates were treated with λ-phosphatase (New England BioLabs) following the manufacturer’s instructions.

### Protein extraction

Cells were lysed on ice for 15 min by adding the appropriate volume of lysis buffer (50mM Tris-HCl pH 7.5, 150mM NaCl, 1mM EDTA, 1mM EGTA, 10mM NaF, 1mM Na3VO4, 0.1% βmercaptoEtOH, 1 tablet of protease inhibitor cocktail, Thermo Scientific).

Cells were harvested by scraping and the lysate was clarified by centrifugation for 15 min at 14000 rcf at 4°C. The supernatants were collected and stored at -80°C.

Bradford reagent (BioRad) has been used as the colorimetric assay for protein measurements.

### Western Blot

20 to 50μg of protein were diluted 1:1 with 2× loading buffer (100mM Tris HCL pH 6.8, 20% glycerol, 4% SDS, 200mM DTT and Bromophenol Blue) and proteins were denatured for 5 mins at 95°C. Samples were resolved on a SDS-PAGE following standard procedures.

Gels were transferred onto Nitrocellulose membrane (GE Healthcare) for 1.5 hrs at 400mA. The membrane was then blocked with 5% Milk in TBS-tween buffer (20mM Tris pH 7.6, 150mM NaCl, 0.1% Tween) for 20 minutes. Membranes were incubated 30 minutes/1 hour or O/N at 4°C in primary antibodies.

Antibodies and dilutions are shown in **Table 2**.

**Table 2.**

| **Antibody** | **Species** | **Dilution** | **Manufactured** |
| --- | --- | --- | --- |
| Cleaved Notch1 | *Rabbit* | 1:1000 | Cell Signalling #4147 |
| β-Actin | *Mouse* | 1:10000 | Proteintech #66009-1-Ig |
| Fbxw7 | *Rabbit* | 1:1000 | Abcam #171961 |
| GFP | *Mouse* | 1:1000 | Cell Signalling #2956 |
| Cdk1 | *Rabbit* | 1:1000 | Cell Signalling #9116 |
| Cdk2 | *Rabbit* | 1:1000 | Cell Signalling #2546 |
| Cdk4 | *Rabbit* | 1:1000 | Santa Cruz #260 |
| Cdk6 | *Rabbit* | 1:1000 | Santa Cruz #177 |
| Cdk8 | *Rabbit* | 1:1000 | Bethyl #A302-501A-T |
| Cyclin A2 | *Mouse* | 1:1000 | Cell Signalling #4656 |
| Cyclin B1 | *Rabbit* | 1:1000 | Cell Signalling #4138 |
| Cyclin C | *Rabbit* | 1:1000 | Bethyl #A301-989A-M |
| Cyclin D1 | *Rabbit* | 1:5000 | Abcam #137875 |
| Cyclin E1 | *Mouse* | 1:1000 | Cell Signalling #4129 |
| Total Histone H3 | *Rabbit* | 1:1000 | Cell Signalling #9715 |
| Phospho Histone H3 | *Rabbit* | 1:1000 | Cell Signalling #3377 |

The membranes were then washed with TBS-Tween and incubated with the appropriate secondary HRP antibody (Cell Signalling, #7074, #7076). After washing, membranes were developed using ECL solution (Pierce).

### Immunoprecipitation

200ug to 1 mg of proteins were incubated O/N at 4°C on rotation with 2 μg of Fbxw7 antibody (Bethyl, A301-721A). Rabbit IgG was used as negative control. Protein G Agarose beads (Cell Signalling #37478) were added for 2 hrs at 4°C on rotation. Precipitates were washed 3 times with cold PBS before adding 2× loading buffer. Samples were denatured for 5 mins at 95°C and then analysed by Western Blot.

Immunoprecipitation using GFP-Trap®_A beads (ChromoTek) was performed according to the manufacturer’s instructions.

### Phos-tag gel

Prior loading into the gels for Phos-tag SDS-PAGE, samples were supplemented with 10 mM MnCl2. Phos-tag SDS-PAGE was carried out as previously described [62, 84]. After electrophoresis, gels were washed three times for 10 min each in the transfer buffer [48 mM Tris/HCl, 39 mM glycine and 20% (v/v) methanol] containing 10 mM EDTA and 0.05% (w/v) SDS, followed by one wash in the transfer buffer containing 0.05% SDS for 10 min. Proteins were transferred and incubated with specific antibodies as previously described.

### Mass Spectrometry

Upon separation by SDS-PAGE on NuPAGE^TM^ 4–12% Bis-Tris protein gels (Thermo Scientific) and staining with Instant Blue (Expedeon), protein bands were excised from gels. Samples were reduced with 10mM DTT at 50°C for 30 min and alkylated with 100mM IAA at room temperature for 20 min in the dark.

Digestion was carried out by adding sequencing grade chymotrypsin (Sigma) at a ratio of 1 to 50 (enzyme to substrate) and incubating O/N at 30°C.

Peptides were extracted with 100% ACN containing 2.5% formic acid and dried in a vacuum centrifuge.

Mass spectrometric analysis was performed by LC-MS-MS using a linear ion trap-orbitrap hybrid mass spectrometer (LTQ-Orbitrap Velos, Thermo Fisher Scientific) and coupled to a Dionex Ultimate 3000 nanoLC system. Peptides were typically injected onto a Thermo Scientific 15cm Easy spray column (Part No. ES800), with a flow of 300 nl/min and eluted over a 40 min linear gradient of 97% solvent A (3% DMSO, 2% Acetonitrile, 0.1% formic acid in H2O) to 35% solvent B (90% acetonitrile, 3% DMSO, 0.08% formic acid in H_2_O). Data files were analysed by Proteome Discoverer 2.0 (Thermo), using Mascot 2.4.1 (www.matrixscience.com), and searching against the Swissprot and MRC-PPU (University of Dundee) databases. Scaffold (www.ProteomeSoftware.com) was also used to examine the Mascot result files. Allowance was made for the following modifications: fixed, Carbamidomethyl (C), and variable, Oxidation (M), Dioxidation (M) and Phosphorylation (S, T and Y). Error tolerances were 10ppm for MS1 and 0.6 Da for MS2. Phospho-sites were assigned according to Proteome Discoverer ptmRS when the phospho-RS site probability was greater than or equal to 90%.

### Kinase assay

Kinase assay was designed and performed by MRC-PPU International centre for Kinase Profiling at University of Dundee (http://www.kinase-screen.mrc.ac.uk/). All assays were carried out using a radioactive (33P-ATP) filter-binding assay.

### Fluorescence-Activated Cell Sorter Analysis

HEK293 cells were harvested, pelleted by gentle centrifugation (450g × 5 min) and washed once in 1 ml of PBS+1% v/v FBS and transferred to FACS tubes (Scientific Laboratory Supplies).

Cells were pelleted again and then fixed in cold 70% ethanol for 30 minutes at room temperature.

Cells number was adjusted to approximately 5×10^5^ cells and then cells were washed twice in PBS+1% v/v FBS.

Cell pellet was then resuspended in 300 μl of staining buffer (50 μg/ml Propidium iodide, 50 μg/ml RNase A in PBS+1% v/v FBS) and incubated for 30 min in the dark at room temperature. FACS analyses were conducted by Flow Cytometry and Cell Sorting facility (University of Dundee).

Briefly, samples were analyzed on a FACS Canto II flow cytometer from Becton Dickinson. Propidium Iodine was detected using 488nm excitation and emission collected at 587 +/™ 40 nm. Data were analyzed using Flowjo software (Flowjo, LLC). Single cells were identified on the basis of PI-A and PI-W measurements and doublets excluded. Cell cycle distribution of the resulting PI-A histograms were determined using the Watson-Pragmatic model.

### Cell cycle synchronization

To synchronize HEK293 cells at G1/S, a double-thymidine block assay was performed. Cells were treated with 2.5 mM of thymidine (Sigma) for 18 hrs, washed twice with PBS, released into fresh media for 9 hrs, treated for a further 14 hrs with 2.5mM of thymidine, then released again into fresh media following two washes in PBS. Cells were then collected for FACS and Western Blot protein analyses at the indicated time points, following protocols already described.

### Mouse Embryo Explant Culture

E10.5 CD1 mice embryos were harvested and explants were prepared as previously described [85]. Each embryo’s posterior part was divided into two halves by cutting along the neural tube. The explants were cultured in hanging drops of culture medium composed of DMEM/F12 (Gibco), 10% FBS, 1% penicillin-streptomycin, 10 ng/ml Fgf2 (PeproTech).

One half was cultured in medium containing Purvalanol B (1 μM) or RO-3306 (10 μM), whereas the control side was cultured in normal medium (DMSO). Both sides were cultured for 4 hr and then analysed for expression of *Lfng* mRNA by in situ hybridization.

For the Western Blot analysis PSM explants from 5 embryos were cultured in DMSO control vehicle and the corresponding contralateral 5 PSM explants cultured in the presence of a small molecule inhibitor.

Experiments were conducted in strict adherence to the Animals (Scientific Procedures) Act of 1986 and UK Home Office Codes of Practice for use of animals in scientific procedures.

### In situ hybridisation of PSM explants using exonic RNA probes

For whole-mount embryos and explants, in situ hybridisation analysis was performed with the use of exonic anti-sense RNA probes as previously described [86].

Samples were imaged using a Leica bright field dissection microscope using Volocity acquisition software. The number of somites present in each explant (identical between explant pairs but variable between embryos) was recorded.

To fix the tissue, samples were incubated for 2 hrs at RT in 4 % PFA/PBS, washed copiously in PBST and transferred into sealed Eppendorf tubes containing 0.01 % sodium azide/PBS for long-term storage at 4°C.

### Statistical Analysis

Unless indicated, Statistical analyses were performed using GraphPad software.

Student t-tests were performed in all the data comparing control to treatment conditions, with P values calculated as * p≤0.05, **p≤0.01 and ***p≤0.001. All experiments have been performed at least three times.

## ACKNOWLEDGEMENTS

We would like to thank Ioanna Mastromina and Michaela Omelkova for reading of the manuscript and constructive feedbacks. Special thanks also go to Genta Ito for assistance with Phostag gel assay. We thank the groups of D. Alessi and S. Rocha for reagents and Laura D’Ignazio for experimental assistance. The authors would like to thank Rachel Toth, Mel Wightman and Tom Macartney (MRC-PPU Reagents and Services) for cloning and Dr David Campbell, Bob Gourlay and Joby Vhargese (MRC-PPU) for Mass Spectrometry analysis. We also thank Professor Daan van Aalten, Professor Victoria Cowling, Dr Francesca Tonelli and Dr Houjiang Zhou for their input.

## COMPETING INTERESTS

I can confirm there are no competing interests for any of the named authors.

## Mathematical model

### 1 NICD - Cell cycle model

To account for the experimental observations of NICD degradation in cell lines, we develop a cell cycle dependent model of NICD production and degradation (see Figure 8A). The position of the *j^th^* cell in the cell cycle is represented by a set of *K* discrete state variables, *X_ij_(t)*, defined such that

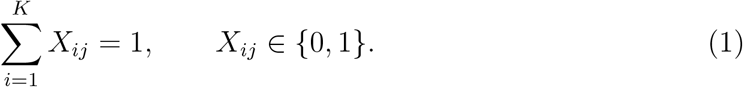

It is assumed that cells transition sequentially between states at rate *r*_1_ and that mitosis is followed by *G*_1_ entry.

The averaged fraction of cells in state *i*, given by

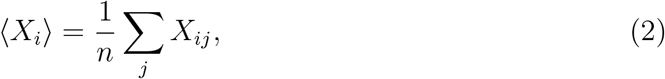

where *n* is the number of cells in the experiment, satisfy

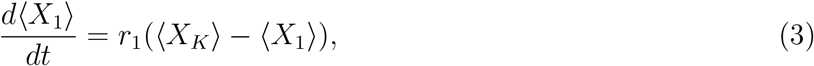

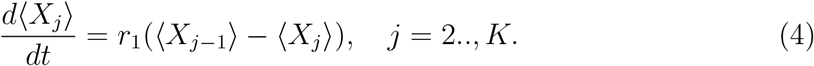

We note that **(author?)** (1) propose a similar model and show that relatively large number of compartments (*K* = 25) compartments can be needed to reproduce experimentally observed variances in cell cycle distribution times.

Letting 〈*X(t)*〉 represent the population average fraction of cells where CDKs are active, then

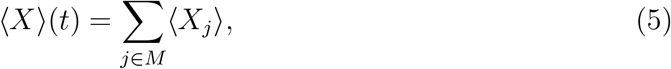

where *M*, the set of CDK active states, has *m* members.

We make the following assumptions about NICD: (i) NICD exists in two states (nonphosphorylated and phosphorylated); (ii) NICD is produced and degraded at constant rates *k*_1_ and *k*_7_, respectively; (iii) NICD is phosphorylated by CDK1 and/or CDK2 at rate *k*_3_;(iv) phosphorylated NICD has a larger degradation rate, *k*_6_, than the background rate, *k*_7_; and (v) dephosphorylation occurs at rate *k*_2_. Letting *N*_1*j*_*(t)* and *N*_2*j*_*(t)* represent the amount of NICD and phospho-NICD in the *j^th^* cell at time *t*, the population averages are defined to be

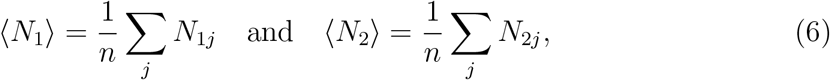

and satisfy

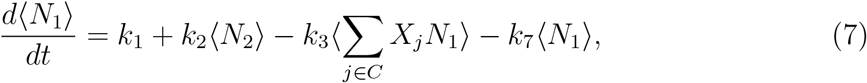

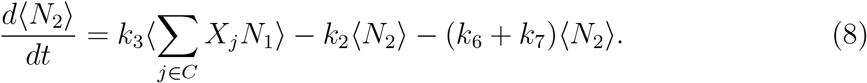

As the cell cycle variables are uncoupled from the NICD dynamics, in a typical cell line experiment each of the *X_j_*’s will tend to the equilibrium value

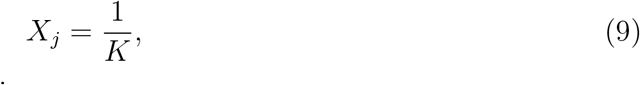

and the fraction of CDK active cells is

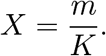

Hence equations (8) reduce to

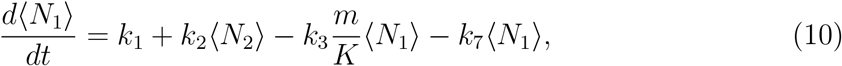

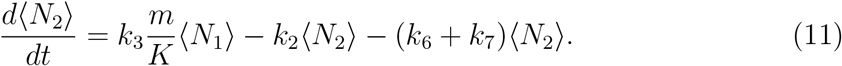

To compute NICD levels equations (11) are solved to steady state 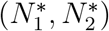 and the total amount of NICD (Figure 8B and 8C) is

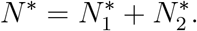

LY treatment experiments, in which the production of NICD is reduced owing to gamma-secretase inhibition, are simulated by setting *k*_1_ = 0. As PurB/Roscovitine is assumed to inhibit CDK2, the effect in the cell cycle synchronised experiments is to reduce the parameter *m*. As MLN treatment inhibits FBXW7 mediated degradation, it is simulated by removing the fast decay mode (i.e. *k*_6_ = 0).

Now consider a situation in which cells are synchronised as a result of double thymidine block. At the point of release the cell cycle distribution is no longer at equilibrium, rather the initial conditions are

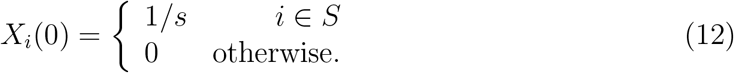

where *S* is a set of states where cells get paused as a result of double thymidine block. After the cells are released from the cell cycle block there are peaks of CDK activity (see Figure 8B) and therefore dynamic levels of NICD.

### 2 Somitogenesis clock and the cell cycle

To explore the coupling between cell cycle mediated NICD degradation and the somitogenesis clock oscillator in PSM tissue, we introduce an additional set of variables, *θ_i_(t)*, that represent the position of the *i^th^* cell in the somitogenesis clock cycle.

As PSM oscillators are coupled via Delta-Notch signalling we consider a phase coupled oscillator model (e.g. 2; 3; 4) of the caudal PSM given

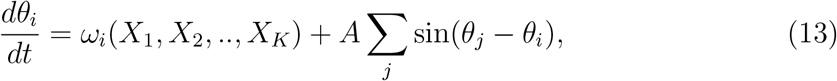

where *ω_i_*, the natural frequency of the *i^th^* cell, is a functional of cell cycle position, the sum is taken over all oscillators and the coupling function represents the effect of Delta-Notch signalling in the caudal PSM.

In a previous model (5) we demonstrated how competition between NICD and Hes7 could yield a phenotype in which increased levels of NICD are correlated with a longer clock period; here we impose this assumption in the phase coupled oscillator model by assuming that the clock frequency is inversely correlated with levels of NICD (Figure 8E). Hence as the cell progresses through the cell cycle, basal levels of NICD fluctuate and modify the natural frequency. We assume that

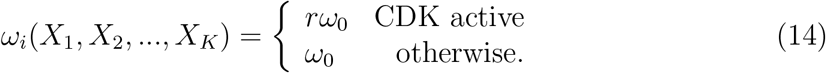

where the parameter *r* represents the degree of acceleration of the clock when CDK is active.

Equations (13) -(14) represent a mathematical model of coupled PSM oscillators whose frequencies are modulated by the cell cycle position. The model allows us to explore whether there is an emergent population-scale period and how it is affected by perturbing the cell cycle. To do this we simulate a population of N cells and find that given adequate oscillator coupling the oscillators synchronise to an emergent frequency which is an average over the individual oscillator frequencies

### 3 Parameter inference

#### 3.1 Cell cycle

The parameters used in the model are presented in Table 2. To recapitulate the cell cycle phase distributions presented in Figure 6 of the Main text we use a 35 state model. The transition rate between compartments, *r*_1_, is defined such that the mean time for a cell to progress from state 1 to is the cell cycle period, *T_C_* = 12h. Based upon the data in Figure 6 in the Main Text we assume the cell cycle times presented in Table 2 (see Figure 1).

**Figure 1:**
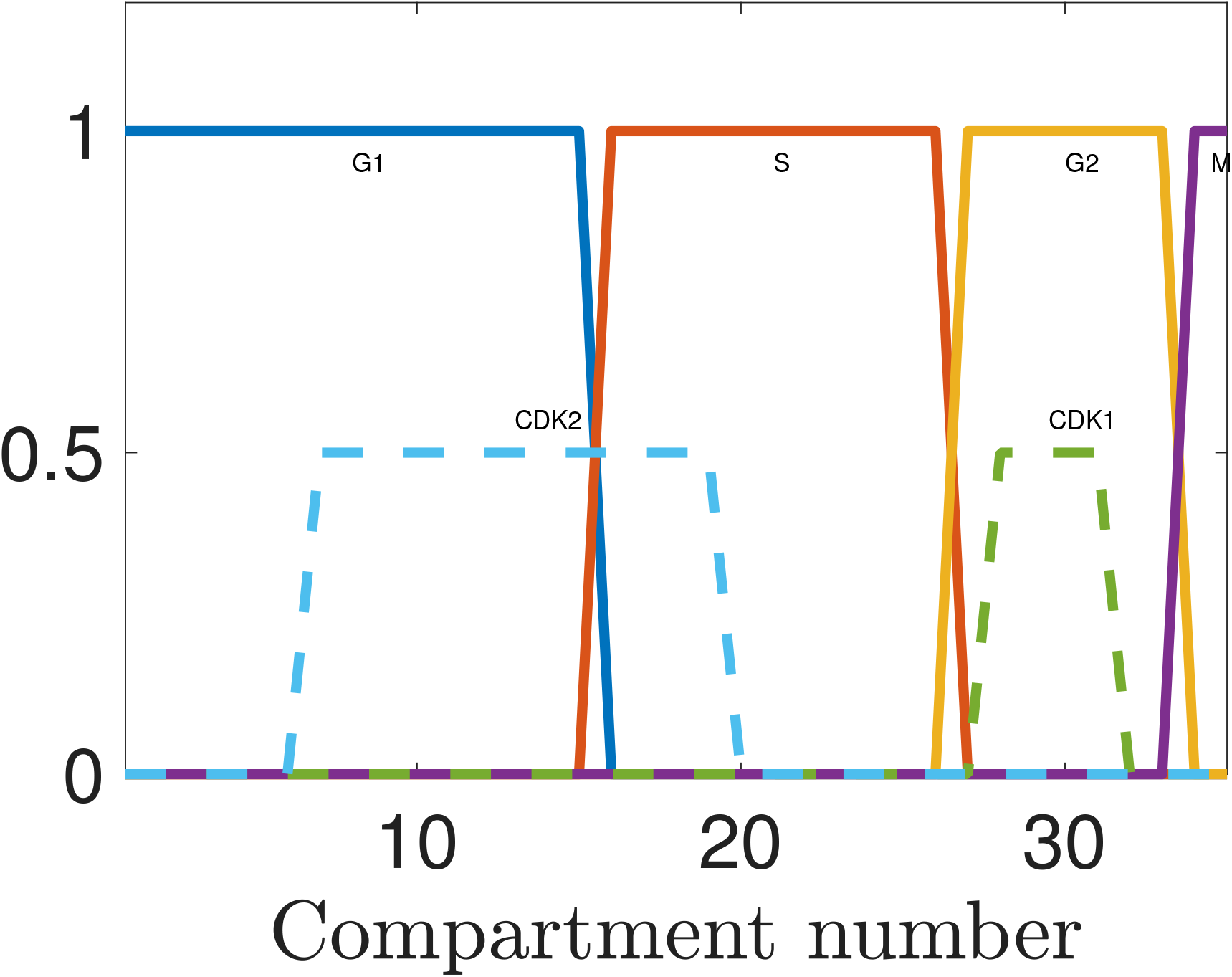
Transition through the cell cycle states (solid lines) and correspondent regions of high CDK2 and CDK 1 activity (dashed lines).

#### 3.2 NICD modelling

The parameter *k*_1_ represents the number of molecules produced per cell per hour. The fast and slow degradation rates are values inferred from **(author?)** (5). To determine the parameters *k*_2_ and *k*_3_ we use the DMSO and MLN treatment Blot data presented in Figure 1 of the Main Text. The measured normalised levels of of non-phosphorylated and phosphorylated forms of NICD in different conditions are given in Table 1. A least squares minimisation algorithm is used to minimise the error between the model and the data in Table 1 for the control and MLN datasets. Hence values for the parameters k_2_ and k_3_ are inferred.

**Table 1:**
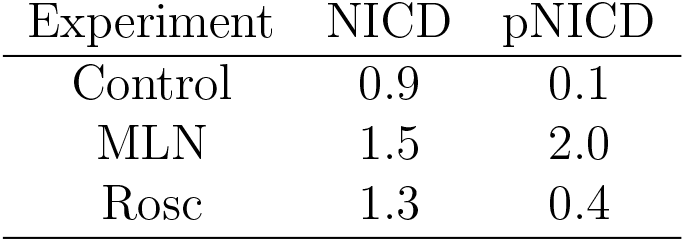
Quantification of Western blots used for inference of parameters *k*_2_ and *k*_3_.

**Table 2:** A table with parameter values.

| Parameter | Value | Unit | Description |
| --- | --- | --- | --- |
| $K$ | 35 | | Number of compartments in cell cycle model |
| $T_C$ | 12.0 | h | Cell cycle period |
| $r_1$ | $\frac{\ln(2)}{KT_C}$ | $\text{h}^{-1}$ | Transfer rate between cell cycle states |
| $k_1$ | 10.0 | $\text{h}^{-1}$ | Rate of NICD production |
| $k_6$ | 12.0 | $\text{h}^{-1}$ | Fbxw7 mediated NICD degradation rate |
| $k_7$ | 1.6 | $\text{h}^{-1}$ | Background NICD degradation rate |
| $k_2$ | 3.1 | $\text{h}^{-1}$ | NICD dephosphorylation rate |
| $k_3$ | 10.0 | $\text{h}^{-1}$ | NICD phosphorylation rate |
| $\omega_0$ | 2.5 | $\text{h}^{-1}$ | somitogenesis clock frequency |
| $A$ | [0,2.0] | $\text{h}^{-1}$ | Oscillator coupling strength |
| $r$ | [1,1.5] | $\text{h}^{-1}$ | Somitogenesis clock acceleration factor |
| $T_{G_1}$ | 4 | h | G1 duration |
| $T_S$ | 3 | h | S duration |
| $T_{G_2}$ | 2 | h | G2 duration |
| $T_M$ | 1 | h | M duration |
| $CDK2_{on}$ | [0.2,0.55] | Nondim | CDK2 active |
| $CDK1_{on}$ | [0.8,0.9] | Nondim | CDK1 active |
| $S$ | [0.4, 0.75] | Nondim | Thymidine block |

#### 3.3 Somitogenesis clock

The somitogenesis clock period in the mouse PSM is approximately two hours. As we do not have direct measurements of coupling strength or acceleration factor we simulate model behaviour over a range the specified parameter ranges.

---

**Supplementary Figure 1.**
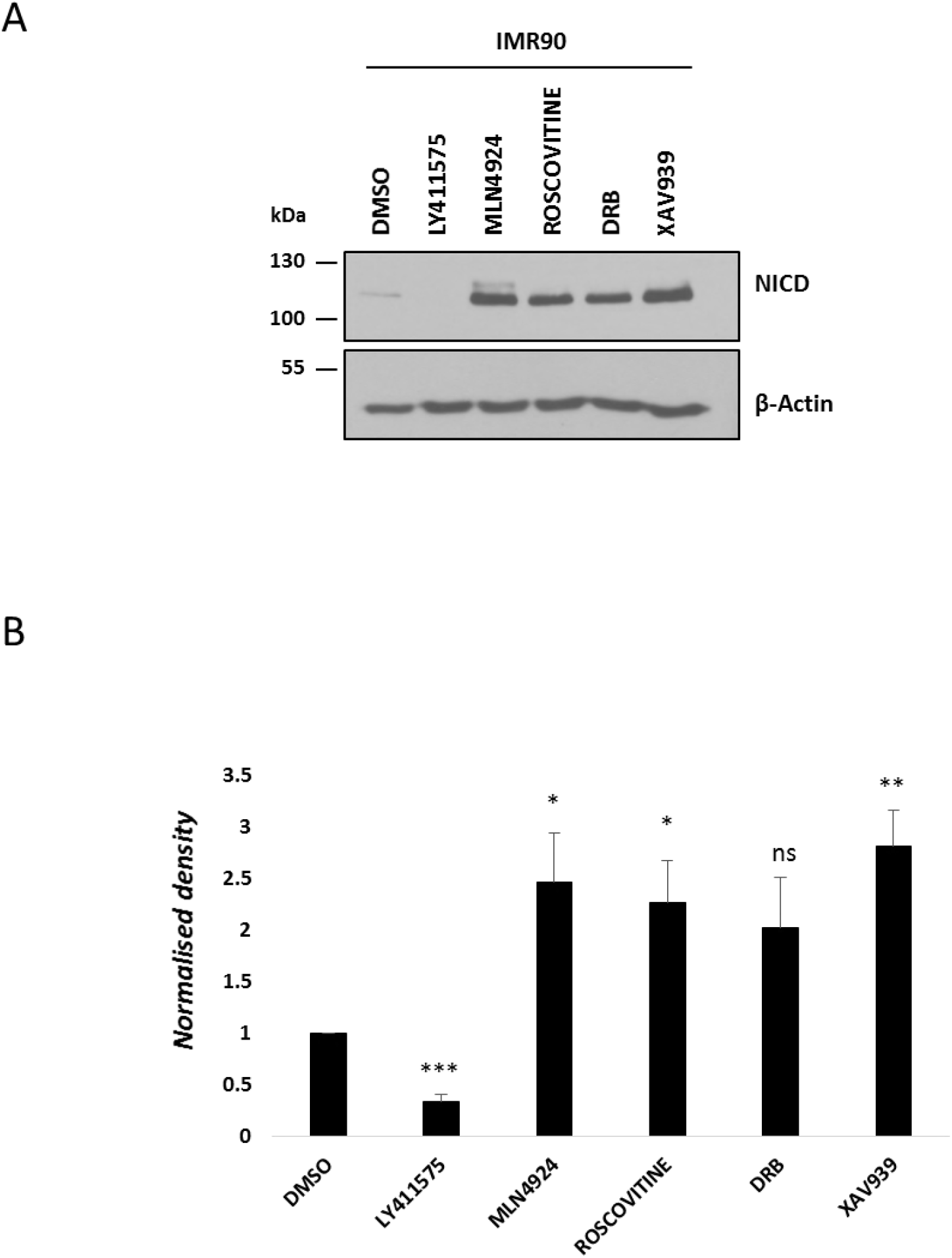
Exposure to Roscovitine, DRB or XAV9393 leads to increased levels of NICD in IMR90 cells. (A) IMR90 cells were treated for 3 hours with 150 nM of LY411575, 1 μM of MLN4924, 10 μM of Roscovitine, 10 μM DRB or 10 μM XAV939. DMSO served as vehicle control. Western blot analysis reveals that NICD levels were increased upon treatment with Roscovitine, DRB, XAV939, and MLN4924. NICD is undetectable following LY411575 treatment. β-Actin served as loading control. (B) Quantification of the density of western blot bands in (A) using ImageJ software. Data are expressed as fold changes compared to DMSO. All data represent the mean ± SEM from three independent experiments. Student’s t-test analysis was performed, with *p≤0.05, **p≤0.01, and ***p≤0.001.

**Supplementary Figure 2.**
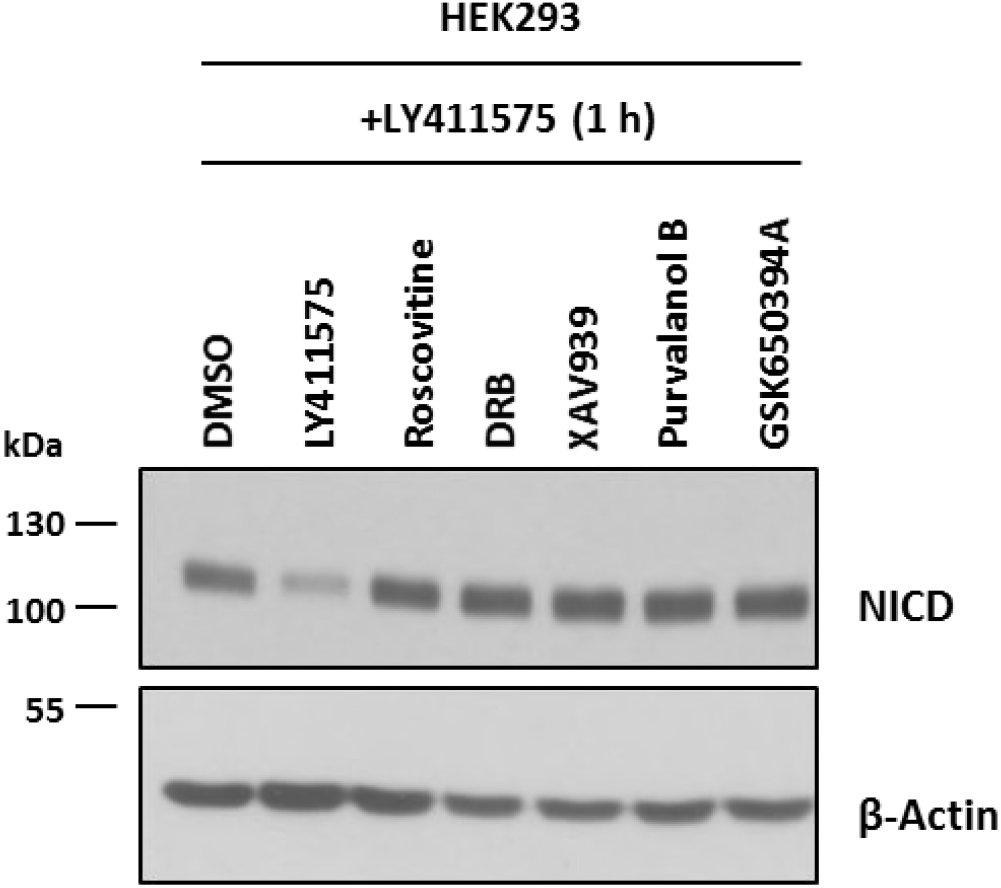
Increase in endogenous NICD levels after exposure to small molecule inhibitors is due to increased stability, not increased NICD production. HEK293 cells were treated for 3 hours with 150 nM of LY411575, 10 μM of Roscovitine, 10 μM of DRB, 10 μM of XAV939, 0.1 μM of Purvalanol B or 10 μM of GSK650394A. DMSO served as vehicle control. 1 hour prior to lysate collection 150 nM of LY411575 was added to DMSO, Roscovitine, DRB, XAV939, Purvalanol B or GSK650394A treated cells to prevent new NICD production. β-Actin served as loading control.

**Supplementary Figure 3.**
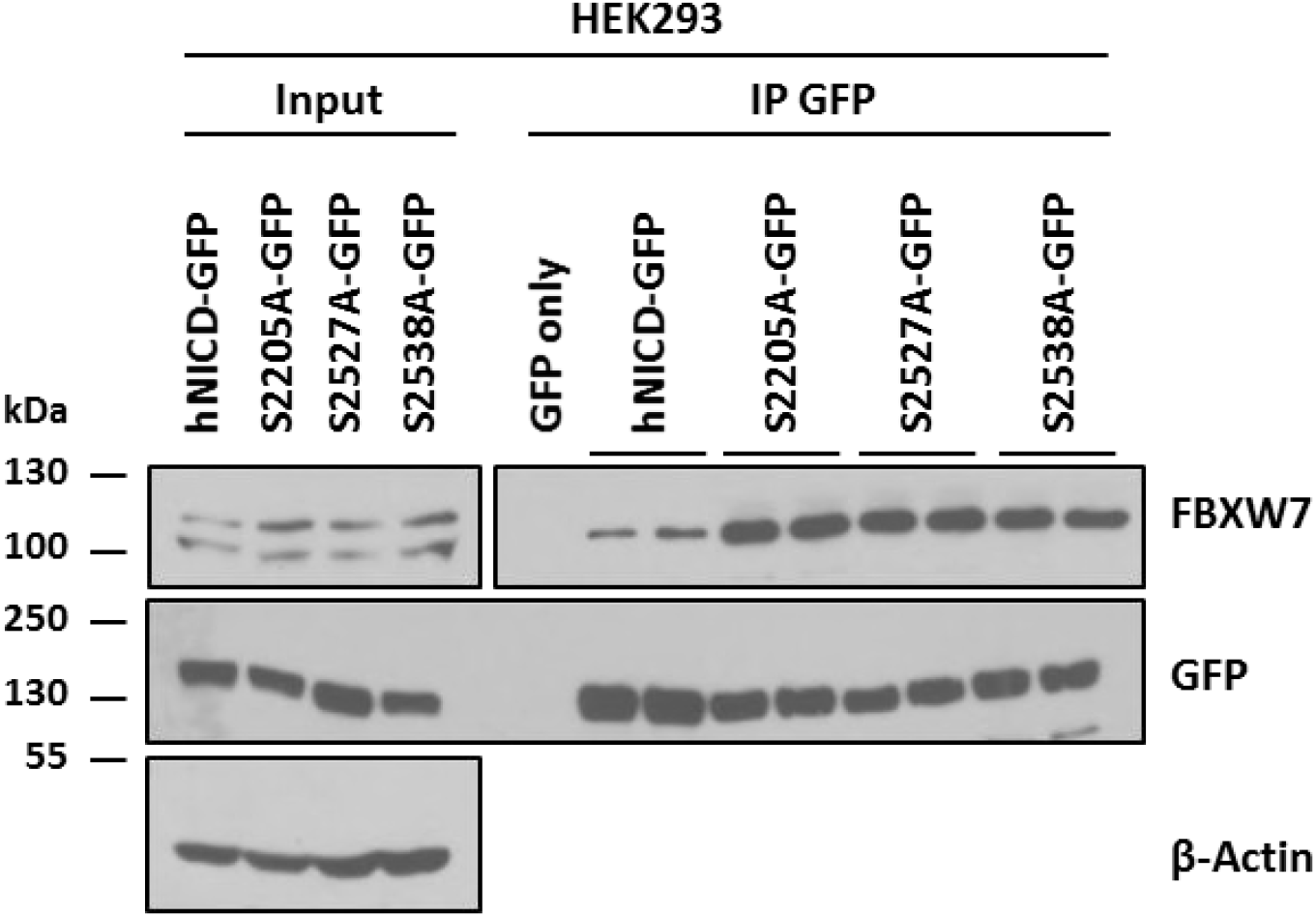
Serine to alanine point mutation on NICD Serine 2205, 2527 and 2538 residues do not change interaction with FBXW7. hNICD-GFP peptides encoding non-phosphorylatable mutations (serine to alanine) were expressed in HEK293 cells. The exogenously expressed protein was subsequently immunoprecipitated with anti-GFP antibody and precipitated material was analysed by western blot using FBXW7 antibody. Wild-type hNICD-GFP and GFP only vectors were included as positive and negative controls, respectively. Western blot using GFP antibody served as immunoprecipitation efficiency control. β-Actin has been used as loading control.

**Supplementary Figure 4.**
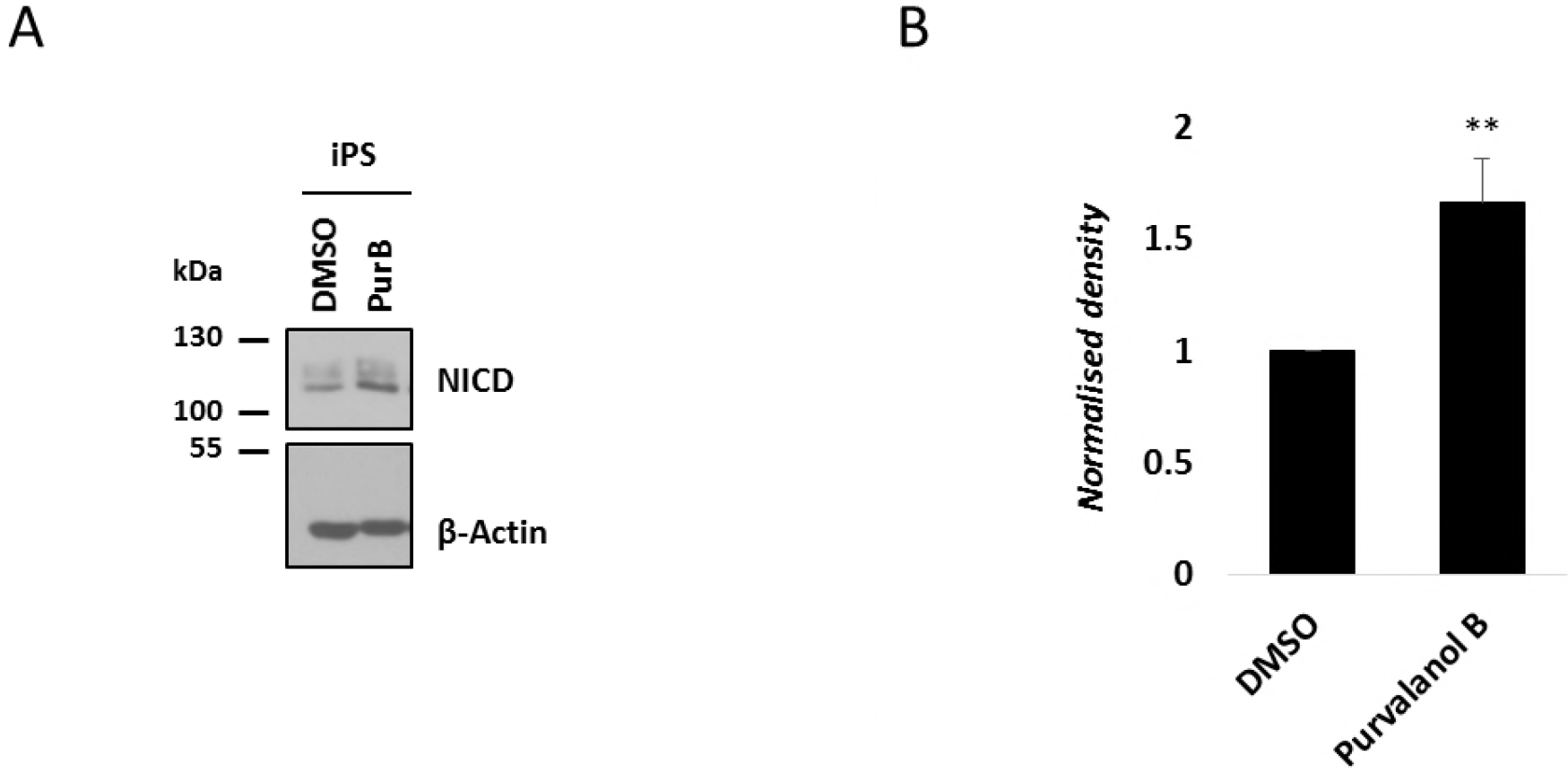
NICD levels increase in iPS cells following treatment with Purvalanol B. (A) iPS cells were treated with 0.1μM of Purvalanol B for 3 hours. Endogenous levels of NICD were detected by western blot. β-Actin has been used as loading control. (B) Quantification of the density of western blot bands in (A) using ImageJ software. Data are expressed as fold changes compared to DMSO. All data represent the mean ± SEM from three independent experiments. Student’s t-test analysis was performed, with **p≤0.01.

